# A hybrid biophysical–neural network model links nanoscale cardiac junctions to tissue conduction

**DOI:** 10.64898/2026.11.16.766810

**Authors:** Ruichen Sui, Xinyu Liu, Nicolae Moise, Dongbin Xiu, Seth H. Weinberg

**Affiliations:** Biophysics Graduate Program, The Ohio State University, Columbus, OH 43210, USA; Davis Heart & Lung Research Institute, The Ohio State University, Columbus, OH 43210, USA; Department of Mathematics, The Ohio State University, Columbus, OH 43210, USA; Department of Biomedical Engineering, The Ohio State University, Columbus, OH 43210, USA

## Abstract

Nanoscale structure of the intercalated disk (ID) governs cardiac conduction through gap junctions and ephaptic interactions and is remodeled in arrhythmogenic disease. Computationally expensive models that resolve ID nanostructure have been confined to cables of tens of cells, such that the role of ID nanostructure on conduction has not been testable. Here, we show that a hybrid biophysical–neural network model removes this limit. A memory-based flow map, trained on finite element method (FEM) simulations of single junctions and short cables, defines surrogates that replace each junction in tissue. The hybrid model reproduces action potentials with 1–2 mV RMSE and conduction velocity restitution across multiple ID geometries with a mean error of 0.24%. Without retraining, surrogate models run in 2-D and 3-D tissues, with over one million junctions, 10^4^–10^5^ times faster than the FEM model and within an order of magnitude of a monodomain model that neglects ID structure. In large tissues with reduced gap-junctional coupling, regional cleft widening blocks conduction, whereas either change alone does not, linking ID remodeling to arrhythmogenic substrate.

## 1 Introduction

Cardiovascular disease is the leading cause of death worldwide, with about 19 million deaths in 2023, one in three of all deaths [1–3]. Sudden cardiac death, mostly from ventricular tachyarrhythmias, accounts for 15–20% of all deaths [4, 5], and atrial fibrillation affects about 60 million people [6, 7]. Many of these arrhythmias arise from slowed or blocked conduction and reentry [8, 9]. Cardiac conduction is governed by the intercalated disk (ID), the myocyte-myocyte junctional region where gap junctions (GJs) reside and mediate electrical coupling.

ID nanoscale structure and molecular composition are remodeled in many arrhythmic diseases. GJs are lost or lateralized in ischemic heart disease, hypertrophy, and heart failure [10]. The perinexal cleft is expanded in patients with atrial fibrillation [11]. Inflammatory vascular leak promotes cleft widening and redistribution of Na^+^ channel clusters, promoting atrial arrhythmias [12]. Changes in ID nanostructure alone alter conduction: conduction velocity depends biphasically on the width of the perinexal cleft, such that widening or narrowing the cleft can slow or accelerate conduction [13, 14], and disrupting cleft adhesion can slow conduction to arrhythmogenic levels [15].

At the ID, Na^+^ channel clusters face each other on adjacent myocyte membranes across a nanometer-scale extracellular cleft, enabling ephaptic coupling in parallel with GJ coupling [16– 18]. ID-localized Na^+^ currents mediate ephaptic coupling, as current into one cell hyperpolarizes the narrow extracellular cleft, which can activate Na^+^ current in the downstream cell. As evidence for this mechanism has come primarily from indirect experiments consistent with ephaptic models [13– 15] and further that representing ID nanostructure adds significant computational cost, standard tissue models, including the monodomain model, minimally represent intercellular coupling by GJs alone. However, recent direct experimental demonstration reported in cardiac cells [18] now shifts the question from whether ephaptic coupling contributes to conduction to how ID nanostructure and its remodeling in disease sets that contribution and its consequences for arrhythmia.

Lumped ephaptic models treat each cleft as one well-mixed extracellular compartment [19, 20]. These models, while more costly than monodomain, are inexpensive enough for tissue simulations and have shown that ephaptic coupling regulates conduction, rescuing conduction in tissue with weak GJ coupling [21] or altering conduction in ischemic border zones [22]. There is increasing evidence that ephaptic coupling also alters repolarization and pro-arrhythmic behavior, such as alternans, early- and delayed afterdepolarizations, and automaticity [23–29]. These lumped models, however, assume a uniform cleft with uniformly distributed channels, and therefore cannot represent the ID nanostructure remodeled in disease.

Spatially resolved models represent the nanoscale distribution of ID membranes and cleft potential, often on finite element method (FEM) grids and mostly with idealized cleft geometries [30–36]. We recently developed an FEM-based ID representation from electron microscopy measurements that captures the structure and ion channel organization within plicate and interplicate regions and cleft electrodiffusion, which we integrated into a short one-dimensional tissue cable [37, 38]. Because every junction carries hundreds of coupled potentials, concentrations, and membrane patches, simulations of these models have been limited to cell pairs or short cables. A 2D tissue large enough to sustain reentry, with 10^5^–10^6^ junctions, is beyond reach. How ID nanostructure, and its remodeling in disease, shapes conduction block in tissue has therefore not been previously testable.

Machine learning offers a path through this bottleneck. Flow map learning (FML) learns the map that advances an observed state by one time step [40], with a memory of past observations when the full state is partially observed [41, 42], and when the system is driven by external inputs [43, 44]. This is the setting of an ID in cardiac tissue: cell interiors interact with a junction only through two cleft currents, which depend on the hidden cleft dynamics through the history of the adjacent cells. Here, we develop a hybrid biophysical–neural network model in which the detailed FEM model [37, 38] is replaced with a memory-based flow map that advances the two cleft currents, computed from their recent history and that of the adjacent cell axial potentials and Na^+^ channel availability (Figure 1). Whereas the monodomain model represents each junction as a fixed conductance, the hybrid model represents it as a learned element, a “smart resistor,” that captures the GJ, ephaptic, capacitive, and ionic currents of a resolved ID. Hybrid models that replace a costly component of a mechanistic model with a learned element have been developed in cardiac modeling, for sarcomere force generation within 3D electromechanics [45] and ion channel kinetics [46]. However, in these studies, the learned component acts within a cell; here, for the first time, the learned component is cell coupling itself, a local element that scales without retraining from a single junction to tissues of any size and topology.

**Figure 1:**
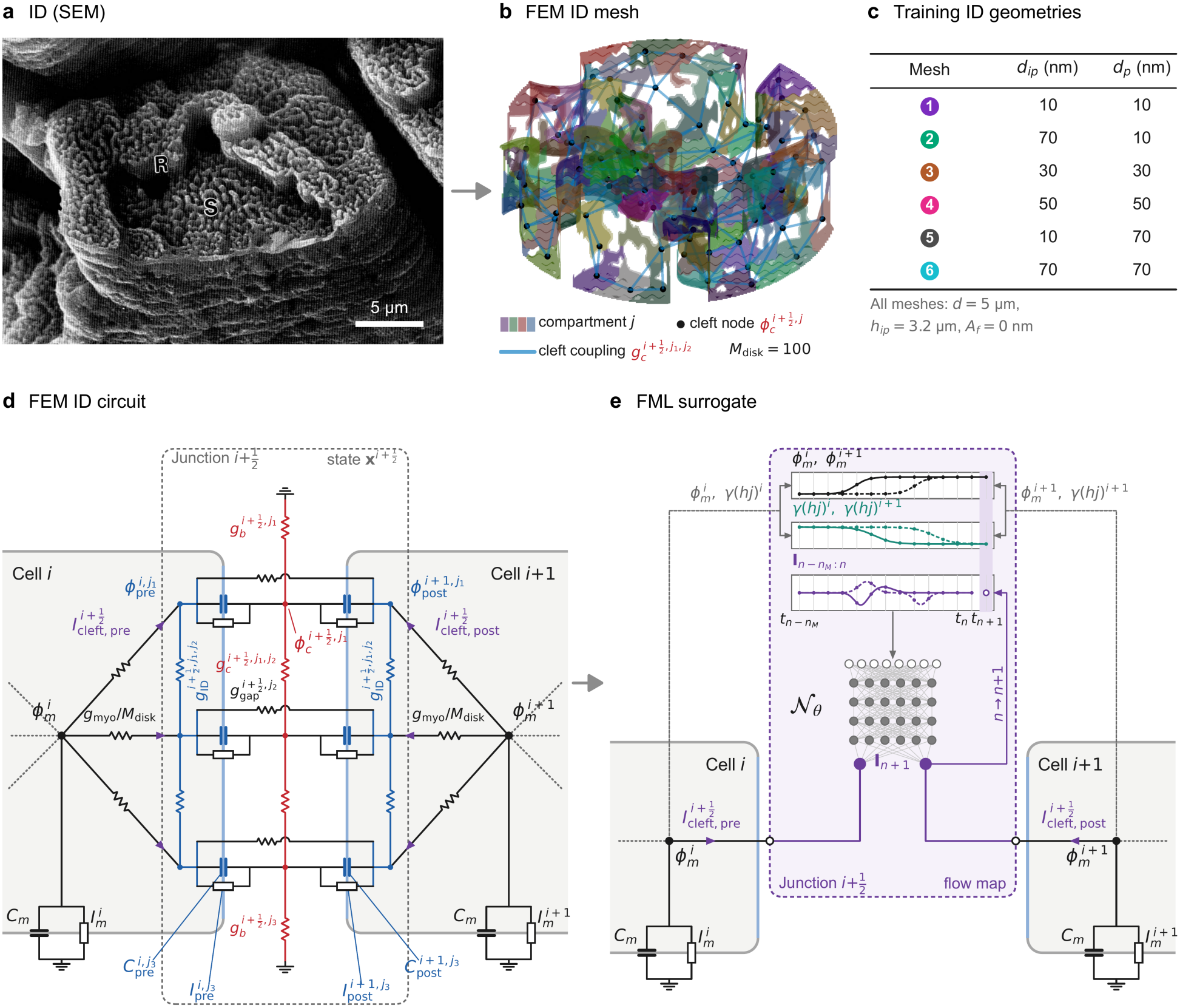
From the ID to the flow-map surrogate in cardiac tissue. (a) Scanning electron micrograph of a ventricular ID of the monkey heart, with steps (S) and risers (R), corresponding to regions conventionally referred to as plicate and interplicate regions; reproduced from [39], Fig. 6 (scale bar from the original magnification). (b) Example FEM ID mesh, partitioned into *M*_disk_ = 100 compartments (one color each); dots, cleft nodes; lines, cleft conductances. (c) ID geometries used for training (meshes 1–6), defined by the interplicate and plicate intermembrane distances *d*_*ip*_ and *d*_*p*_; all meshes: *d* = 5 µm, *h*_*ip*_ = 3.2 µm and *A*_*f*_ = 0 nm. (d) FEM ID circuit of junction 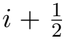 between cells *i* and *i* + 1, illustrating three of the *M*_disk_ compartments (*j*_1_, *j*_2_, *j*_3_). The axial node of each cell is coupled to every compartment through a resistor with conductance *g*_myo_*/M*_disk_. Pre- and post-junctional ID patches (blue) face a common cleft node (red), which is coupled to the other cleft nodes (and potentially to the grounded bulk). The cells see the junction state 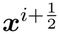 (dashed box) only through the cleft currents 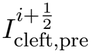 and 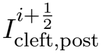 (purple arrows). (b,d) Adapted from [38]. (e) FML surrogate. The junction is replaced by the flow map ***N*** _*ω*_ (Equation (9)), which maps the histories of 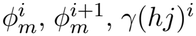, *γ*(*hj*)^*i*^ and *γ*(*hj*)^*i*+1^ (cell *i*, solid; cell *i* + 1, dashed) up to the target time *t*_*n*+1_ (shaded), and of the cleft currents 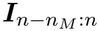, to ***I***_*n*+1_. The cleft current outputs are supplied directly to the cell axial membrane and appended to the current history, such that each cell is connected to each adjacent junction through a single port (dotted lines).

Here, we show that this hybrid model reproduces the FEM model across coupling regimes, ID geometries, and pacing rates, and that networks trained on cables run without retraining in heterogeneous cables and in 2D and 3D tissue with up to 1.2 million junctions. The hybrid surrogate model is 10^4^–10^5^ times faster than the FEM model and within an order of magnitude of a monodomain model that omits the ID. One beat of a 3D tissue block, requiring an estimated 2.5 years with the FEM model, can be simulated in 3 minutes. These simulations show that regional cleft widening can produce conduction block, linking nanoscale ID remodeling to arrhythmogenic substrate.

## 2 Results

We first identify the most accurate network configuration (Section 2.1) and compare the surrogate with the FEM model across GJ coupling, ID geometries, and pacing rates (Section 2.3) and in a heterogeneous cable (Section 2.4). We then simulate large 2D and 3D tissues with up to 1.2 million junctions, compare computational costs, and demonstrate how regional heterogeneity in ID nanostructure alters conduction (Section 2.5). Unless stated otherwise, simulations use an implicit integration scheme with *n*_sub_ = 4 and the five-member ensemble. Protocols and error measures are given in Section 4.6.

### 2.1 Memory and channel availability improve surrogate accuracy

We determine the relationship between the accuracy of the surrogate model and two aspects of the network configuration, the duration of memory *T*_*M*_ and the network source (***u***, Eqn. (5)), specifically the inclusion of the scaled Na^+^ channel availability *γ*(*hj*) of the two cells, in addition to the cell axial potentials *ϕ*_*m*_. We measure *ϕ*_*m*_ RMSE in a 50-cell cable, with ID properties from mesh 1 (*d*_*ip*_ = *d*_*p*_ = 10 nm), for both GJ coupling strengths and BCLs of 300 and 200 ms (Figure 2; traces in Supplementary Figures S1 and S2).

**Figure 2:**
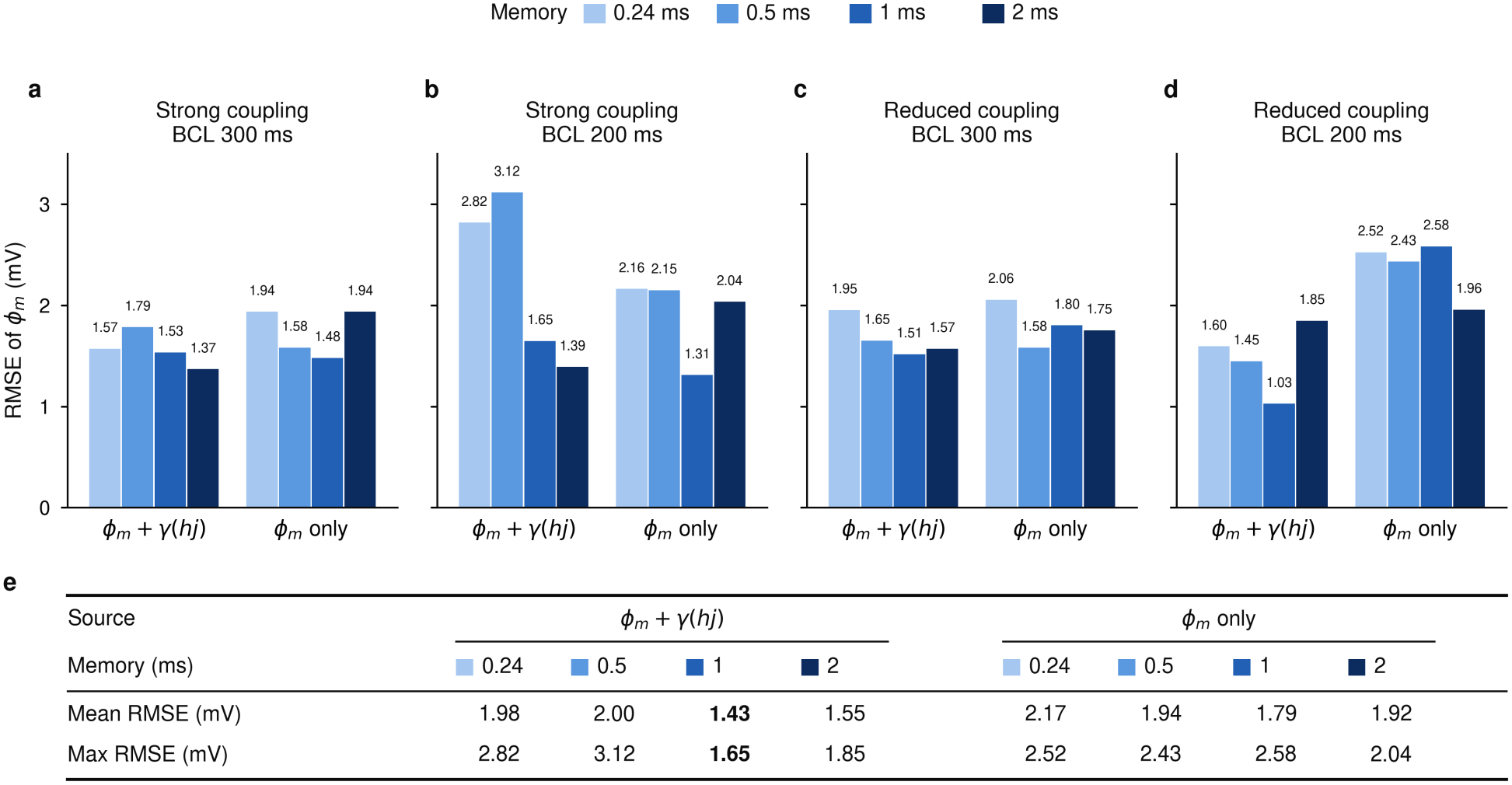
Network configuration choice improves reconstruction. RMSE of the axial potential *ϕ*_*m*_ over all 50 cells and the ten beats, from the surrogate model against the FEM reference for mesh 1. The surrogates use the full source ***u*** of Equation (5), labeled *ϕ*_*m*_ + *γ*(*hj*), or the axial potentials only, labeled *ϕ*_*m*_ only, each with the memories *T*_*M*_ = 0.24, 0.5, 1, and 2 ms. (a–d) Strong (a,b) and reduced (c,d) GJ coupling at BCL 300 ms (a,c) and 200 ms (b,d). (e) Mean and maximum of the RMSE over the four cases; the smallest values are in bold.

We find that all configurations reproduce the action potential upstroke and cleft currents and differ mainly in repolarization. Including Na^+^ availability, increasing the memory up to 1 ms improves accuracy in all four cases, with the largest improvement for strong coupling at BCL of 200 ms. Increasing the memory further to 2 ms does not systematically reduce the error and in some cases increases it. Error for a 2 ms memory is lower with strong GJ coupling but increases with reduced coupling at BCL of 200 ms (1.85 vs. 1.03 mV). At 1 ms memory, the source *ϕ*_*m*_ case is slightly more accurate with strong GJ coupling, but with reduced GJ coupling, error is more than doubled. Across all conditions, 1 ms memory duration and inclusion of both the axial potentials and Na^+^ channel availability has the smallest mean and smallest maximum RMSE, compared with every other configuration (Figure 2e). A memory duration of 1 ms also halves the network input relative to 2 ms (310 instead of 610 values), which in turn reduces the arithmetic of each network evaluation by about a third. Thus, we use this configuration for all results that follow.

### 2.2 Surrogate model reproduces action potential characteristics

Figure 3 illustrates conduction in the cable with mesh 1 paced at BCLs of 300 and 200 ms for both strong and reduced GJ coupling. In all four cases, *ϕ*_*m*_ and the cleft currents along the cable from the surrogate model closely agree with the FEM reference (Figure 3a). The closed-loop simulation is stable, with no drift of the resting potential or of the plateau between beats. *ω*_*m*_ RMSE is between 1-2 mV for all cases, with close matching of the action potential (AP) upstrokes (Figure 3b). Conduction velocity (CV) of the last beat is within 0.3% of the reference (BCL of 300 ms: strong GJ coupling 36.98 (surrogate) vs. 37.07 cm s^*−*1^ (FEM); reduced GJ coupling 18.27 (surrogate) vs. 18.31 cm s^*−*1^ (FEM)). We observe minimal differences throughout the AP, with the largest differences primarily confined to late repolarization, largest for reduced GJ coupling at BCL of 300 ms.

**Figure 3:**
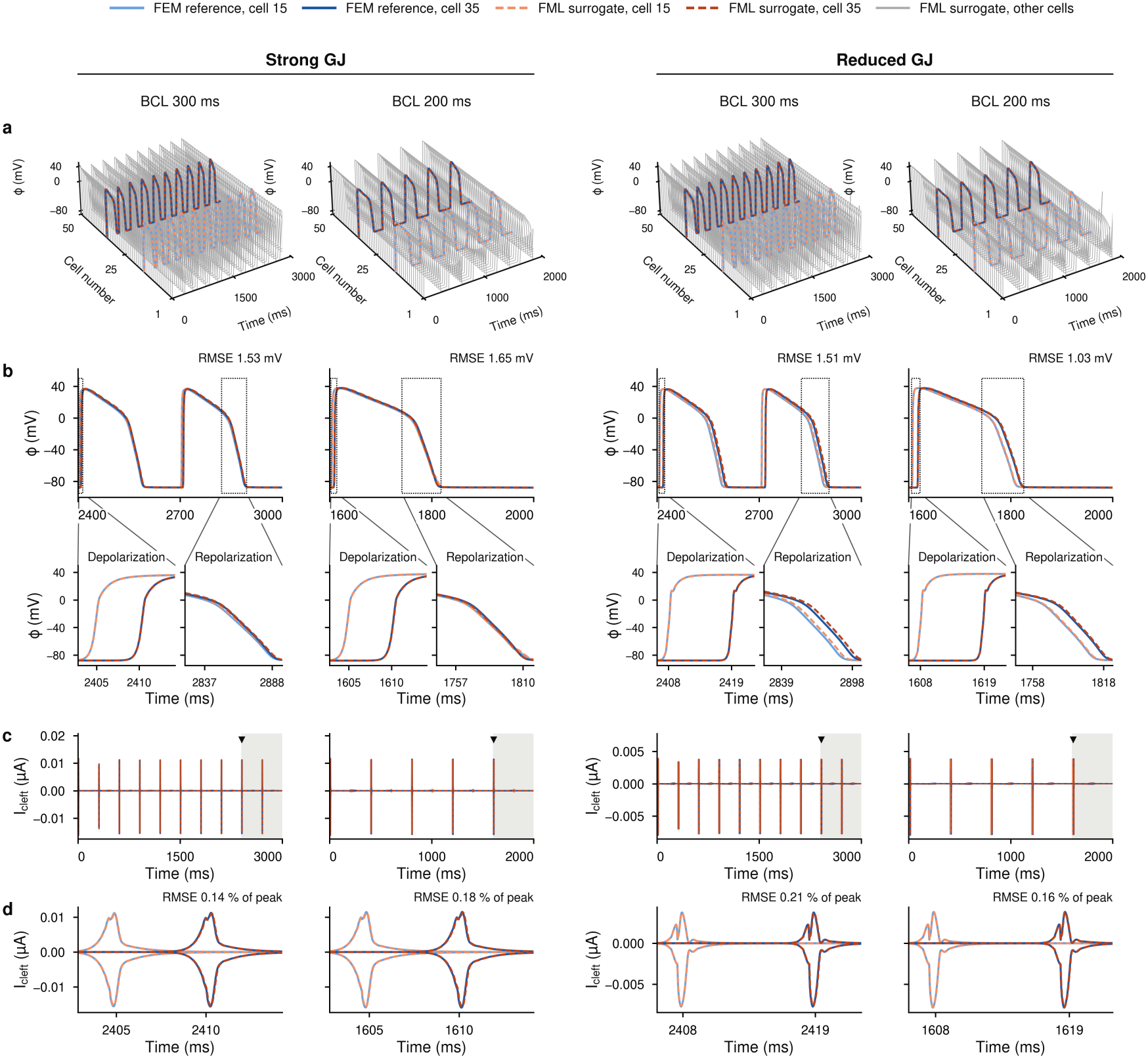
Closed-loop simulation of a 50-cell cable with mesh 1 (*d*_*p*_ = *d*_*ip*_ = 10 nm, Figure 1c), for strong (left) and reduced (right) GJ coupling and periodic pacing at BCLs of 300 ms and 200 ms. (a) Axial potentials of all cells; cells 15 and 35 are highlighted. (b) Last two beats of cells 15 and 35, with enlarged views of depolarization and repolarization. (c) Cleft currents over the whole run at junctions 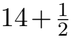 (light, between cells 14 and 15) and 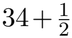 (dark, between cells 34 and 35). Each junction has two curves, the current into the pre-junctional (upstream) side, positive, and into the post-junctional (downstream) side, negative. The shaded window is shown in (b), and the triangle marks the beat expanded in (d). (d) Cleft currents during the first upstroke of the last two beats.

The cleft currents, the quantity the surrogate model predicts, are primarily nonzero during the AP upstroke (Figure 3c). For strong GJ coupling, we find that the cleft current is a single transient, positive into the upstream and negative into the downstream cell. For reduced GJ coupling, the cleft current peak magnitude is reduced, as expected, and exhibits a broader time course (Figure 3d). Note that pre- and post-junctional currents differ in amplitude and shape and are not symmetric (unlike the equal and opposite currents of an ohmic resistor), consistent with their independent predictions (Section 4.4). Throughout the cable, the surrogate model reproduces cleft current amplitude, timing, and multi-peak structure, with an RMSE of 0.14 %–0.21 % of the peak current.

### 2.3 Surrogate model captures conduction velocity restitution

We next investigate if the surrogate model can accurately reproduce the rate dependence of CV across ID geometries. Conduction slows as the pacing rate increases (i.e., BCL decreases), a property known as CV restitution. CV restitution, together with the restitution of the action potential duration, are key properties that shape discordant alternans, conduction block, and the induction of reentry [47–49]. We paced the cable with each of the six ID geometries for both strong and reduced GJ coupling (Figure 1c) at BCLs decreasing from 1000 ms to conduction failure (Figure 4). CV restitution of the surrogate agrees with that of the FEM reference in all twelve cases. Over the 178 measurement points, the mean absolute CV error is 0.24 % and the largest 1.18 % (reduced coupling, mesh 3, BCL 230 ms). The errors are largest for the shortest BCLs, where the CV falls steeply, and remain below 0.5% for BCLs of 400 ms and longer. In every case, the surrogate also reproduces the BCL for which which conduction no longer follows every stimulus.

**Figure 4:**
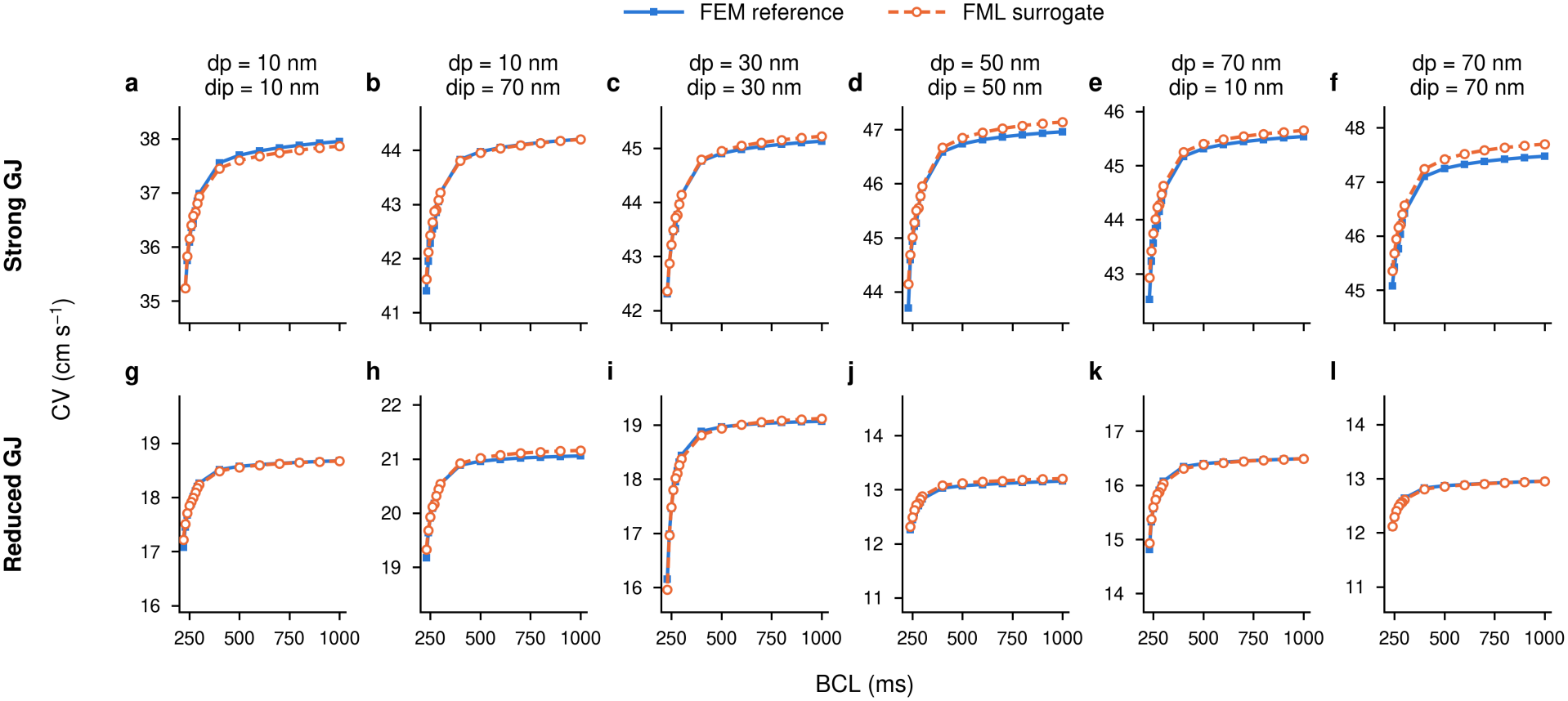
CV restitution of the 50-cell cable for the six ID geometries (columns, meshes 1–6) with strong (a–f) and reduced (g–l) gap-junctional coupling. Each point is the CV at the given BCL, for the FEM reference (solid) and the surrogate (dashed, five-member ensemble). The curves run from BCL 1000 ms down to the last BCL for which beats conduct in a 1:1 pattern.

Critically, the surrogate models capture the dependence on ID properties, specifically the plicate (*d*_*p*_) and interplicate (*d*_*ip*_) widths, as demonstrated in the FEM model. Specifically, we previously showed that the influence of the plicate and interplicate widths can speed or slow conduction, depending on the GJ coupling [38], and these relationships are also captured by the surrogate:

For strong GJ coupling, widening either the plicate or interplicate speeds up conduction, while for reduced GJ coupling, the same widening slows conduction. For reduced GJ coupling, the dependence on ID geometry is more complicated, with changes in CV as *d*_*p*_ varies in turn depending on *d*_*ip*_, and vice versa, and the surrogate reproduces all of these changes with high accuracy.

### 2.4 Accurate prediction of conduction in a heterogeneous cable

In physiological cardiac tissue, ID geometry will vary from one junction to the next. From a model perspective, this corresponds to each junction having neighbors of a different geometry and thus presents the challenge of receiving inputs unlike those of its training data, which are generated from homogeneous cables (Section 4.4). To test reproduction accuracy in this more realistic setting, we simulate a 50-cell cable for which the mesh of each of the 49 junctions is drawn at random from the six meshes (Section 4.6; Figure 5, top). As before, we compare the surrogate model with a comparable cable with the ID represented by the FEM model for both strong and reduced GJ coupling.

**Figure 5:**
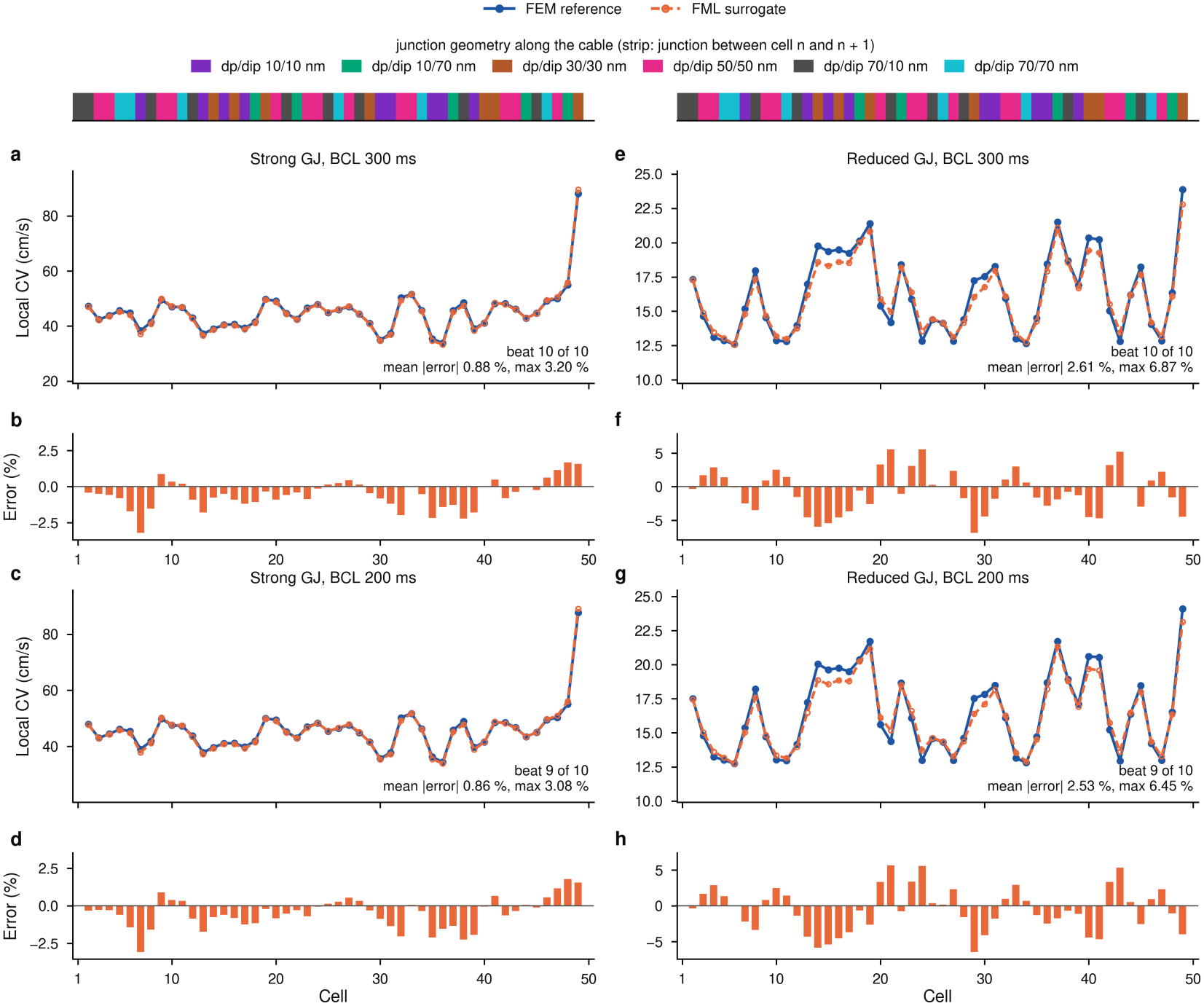
Local CV along the 50-cell cable with heterogeneous ID geometry. CV shown for the last propagated beat: beat 10 at BCL 300 ms and beat 9 at BCL 200 ms. (a,c,e,g) Local CV of the FEM reference and the surrogate at every cell for strong (a,c) and reduced (e,g) gap-junctional coupling; the mean and maximum absolute relative errors over the cable are given in each panel. (b,d,f,h) Relative error of the surrogate at every cell. The strips at the top show the junction geometry along the cable.

For BCL 300 ms, every beat propagates along the cable, while for 200 ms, only the odd beats propagate, and the surrogate reproduces this 2:1 pattern exactly (Supplementary Figure S3). The RMSE of the potential is 2.08 mV–3.27 mV, somewhat larger than in the homogeneous cable and again confined to late repolarization, and that of the cleft currents is 0.3 %–0.8% of the peak current. The local CV along the cable (Equation (17)) varies strongly from cell to cell, between about 35 and 50 cm s^*−*1^ with strong GJ coupling and between about 12.5 and 22 cm s^*−*1^ with reduced GJ coupling. The surrogate model follows the reference FEM model CV at every cell (Figure 5), with mean absolute error of 0.86 %–0.88 % with strong GJ coupling (maximum 3.20 %) and 2.53 %–2.61 % with reduced GJ coupling (maximum 6.87 %). The error does not exhibit a bias in a particular direction, changing sign along the cable and does not increase with the distance from the stimulus, demonstrating that error does not accumulate.

### 2.5 Conduction block in large two- and three-dimensional tissues

Finally, we demonstrate the feasibility of simulating larger and higher-dimensional tissues that incorporate the effects of ID nanostructure. We first validate the surrogate model on a 20 ×20 2D sheet, small enough to simulate with the FEM model, for both strong and reduced GJ coupling, and we find close matching of the surrogate activation time and action potentials with the FEM reference, with *ϕ*_*m*_ RMSE between 5-10 mV (Supplementary Figure S4). For the larger tissues below, the FEM model cannot be run on our workstation, due to memory limitations and prohibitively high computational cost.

We quantify the computational costs for tissues of increasing dimension and size, comparing the FEM model with the surrogate model for different implementations and integration schemes (Table 1; hardware in Supplementary Table S2). The FEM model costs about 0.3 s per junction and simulated millisecond, whereas the surrogate evaluates the networks of all junctions together, such that the computational cost efficiency increases with tissue size: about 260 times in the cable, and with the implicit scheme on the GPU, 5.5 × 10^4^ times in the 2D sheet, 4.5 ×10^4^ times in a 20 ×20 ×8 3D block and an estimated 2.5 ×10^5^ times in the largest 3D block (0.8–1.5 ×10^4^ times on the CPU). One beat of the 100 ×100× 40 3D tissue block at a BCL of 205 ms would require an estimated two and a half years with the FEM model and takes about 3 min with the surrogate (explicit scheme) on one GPU.

**Table 1:** Wall time of the time loop for 1 ms of simulated time, per step, and speed-up relative to the FEM model with Δ*t* = 0.001 ms on the same tissue. Per step: one step Δ*t* of the FEM model, one macro step Δ*t* of the surrogate. Explicit: explicit; implicit: implicit (Section 4.5); settings in Section 4.6. The 100 ×100× 40 block does not fit into memory with the FEM model: its FEM time and speed-ups are estimates (est.), scaled from the 20 ×20 ×8 block by the number of junctions. Monodomain: the same tissue with every junction replaced by an ohmic gap junction, with the same implementation, step and scheme; ratio: wall time of the surrogate over that of the monodomain model.

|  |  |  |  |  | Monodomain |  |
|---|---|---|---|---|---|---|
| Tissue | Code | 1 ms simulated | Per step | Speed-up | 1 ms | Ratio |
| 1-D cable, 50 cells | MATLAB FEM | 15.40 s | 15.40 ms | 1 | – | – |
|  | JAX, CPU, implicit | 0.05863 s | 1.173 ms | 263× | – | – |
|  | JAX, CPU, explicit | 0.02848 s | 0.5697 ms | 541× | – | – |
|  | JAX, GPU, implicit | 0.1801 s | 3.601 ms | 86× | – | – |
|  | JAX, GPU, explicit | 0.08155 s | 1.631 ms | 189× | – | – |
| 2-D sheet, 100 × 100 cells | MATLAB FEM | 6550 s | 6550 ms | 1 | – | – |
|  | JAX, CPU, implicit | 0.4337 s | 8.674 ms | 15 103× | 0.04229 s | 10.3× |
|  | JAX, CPU, explicit | 0.2819 s | 5.639 ms | 23 233× | 0.03783 s | 7.5× |
|  | JAX, GPU, implicit | 0.1197 s | 2.393 ms | 54 739× | 0.04318 s | 2.8× |
|  | JAX, GPU, explicit | 0.07031 s | 1.406 ms | 93 153× | 0.006862 s | 10.2× |
| 3-D block, 20 × 20 × 8 cells | MATLAB FEM | 2903 s | 2903 ms | 1 | – | – |
|  | JAX, CPU, implicit | 0.2019 s | 4.039 ms | 14 374× | 0.01369 s | 14.7× |
|  | JAX, CPU, explicit | 0.1168 s | 2.336 ms | 24 852× | 0.01251 s | 9.3× |
|  | JAX, GPU, implicit | 0.06436 s | 1.287 ms | 45 099× | 0.02736 s | 2.4× |
|  | JAX, GPU, explicit | 0.04424 s | 0.8847 ms | 65 617× | 0.005222 s | 8.5× |
|  | CUDA engine, implicit | 0.1663 s | 3.326 ms | 17 454× | 0.06337 s | 2.6× |
|  | CUDA engine, explicit | 0.09167 s | 1.833 ms | 31 663× | 0.007702 s | 11.9× |
| 3-D block, 100 × 100 × 40 cells | MATLAB FEM (est.) | 3.86 × 10<sup>5</sup> s | 386 s | 1 | – | – |
|  | JAX, CPU, implicit | 46.86 s | 937.2 ms | 8250× | 3.348 s | 14.0× |
|  | JAX, CPU, explicit | 26.42 s | 528.3 ms | 14 600× | 2.762 s | 9.6× |
|  | JAX, GPU, implicit | 1.570 s | 31.40 ms | 246 000× | 0.2322 s | 6.8× |
|  | JAX, GPU, explicit | 0.9913 s | 19.83 ms | 390 000× | 0.1874 s | 5.3× |
|  | CUDA engine, implicit | 1.460 s | 29.20 ms | 265 000× | 0.2820 s | 5.2× |
|  | CUDA engine, explicit | 0.9488 s | 18.98 ms | 407 000× | 0.2172 s | 4.4× |

We also evaluate the surrogate model against a comparable monodomain model of the same tissues, in which every junction is reduced to an ohmic gap junction without detailed ID and cleft representation. This simpler representation ranges from 2.4–14.7 times faster than the surrogate with the same implementation, hardware, and scheme (Table 1). Our hybrid surrogate thus brings tissue simulations that resolve the ID nanostructure to within about one order of magnitude of the cost of the monodomain model.

Our final results illustrate how regional heterogeneity in ID nanostructures can drive irregular conduction and conduction block. We simulated a sheet of 100 ×100 cells (19,800 junctions) and a block of 100 ×100 ×40 cells (1,182,000 junctions). For both cases, GJ coupling was uniformly reduced throughout the tissue with nominal ID properties (mesh 1, *d*_*p*_ = *d*_*ip*_ = 10 nm) in most regions and disrupted clefts (mesh 4, *d*_*p*_ = *d*_*ip*_ = 50 nm) in a central disk or ellipsoid (Figure 6; Supplementary Videos 1 and 2). In the 2D sheet, the first beat of a propagating planar front is delayed in the heterogeneous region due to slower conduction (Figure 6a). As the BCL is gradually decreased, the region is activated progressively later, leading to conduction block in this region: at a BCL of 206 ms, the region conducts only every second beat, while the surrounding tissue conducts every beat (Figure 6b,c). In the 3D block, conduction is similarly slowed through the heterogeneous region (Figure 6d). On the ninth beat, the wave blocks at the center of the ellipsoid (Figure 6e,f), propagates around the heterogeneous region and enters its distal half from behind, while the surrounding tissue conducts the beat as a plane wave; the tenth beat conducts again. In both cases, slowed conduction in the wide-cleft region results in extended refractoriness that blocks conduction on the subsequent beat. In contrast, in two control tissues, one with reduced GJ coupling but without the wide-cleft region and one with a wide-cleft region but with strong GJ coupling, every beat conducts (Supplementary Figure S5), demonstrating that the combination of disrupted IDs and reduced GJ coupling drives conduction block.

**Figure 6:**
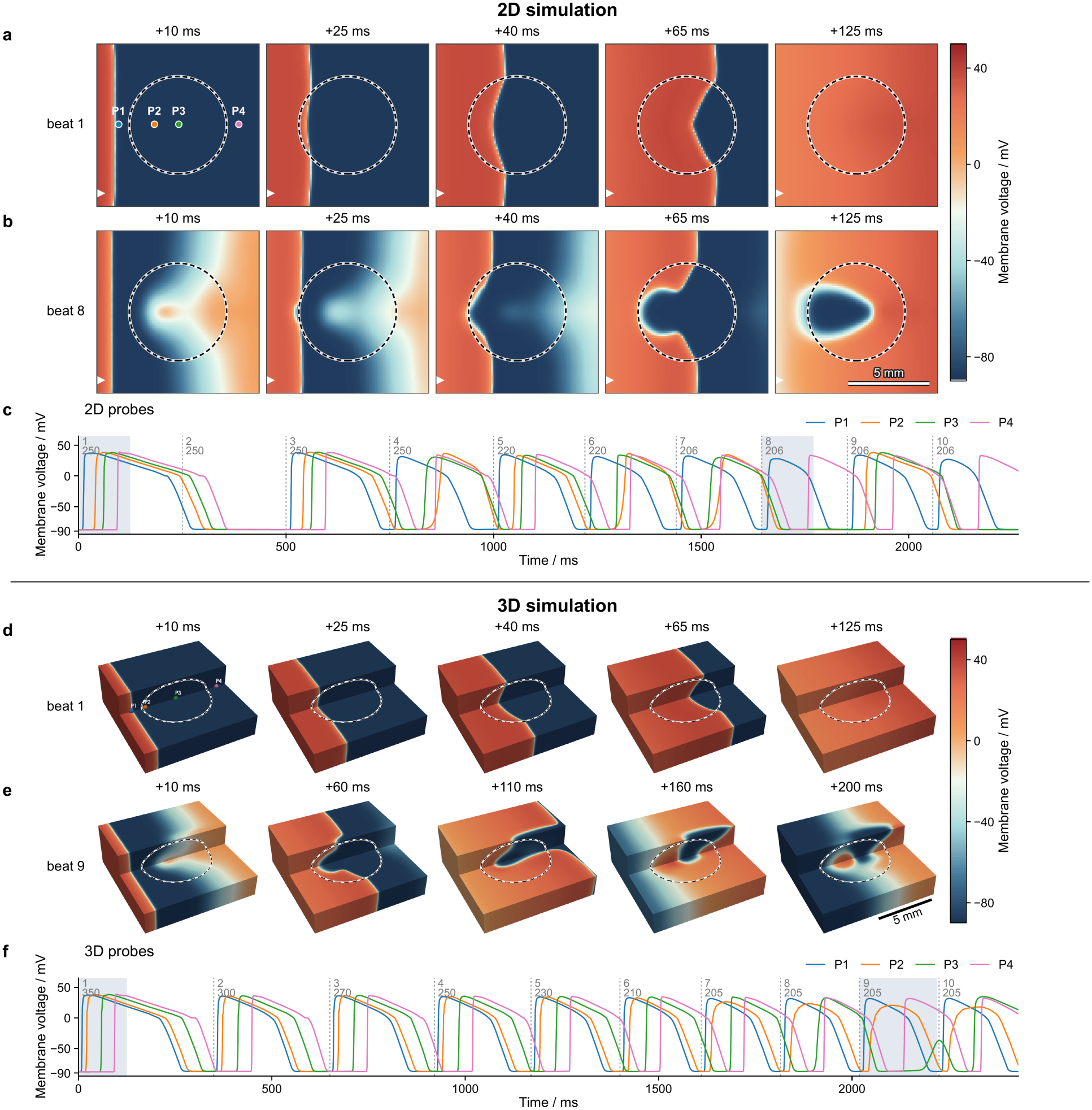
Conduction block in 2-D and 3-D tissues with regional heterogeneity and reduced GJ coupling. Junctions inside a central region (dashed circle in 2-D, black line on the visible faces in 3-D) use mesh 4 (*d*_*p*_ = *d*_*ip*_ = 50 nm), all others mesh 1 (*d*_*p*_ = *d*_*ip*_ = 10 nm). 2D simulation: (a,b) Sheet of 100 × 100 cells with a disk of radius 30 cells, paced at the left edge (arrowhead) with the cycle length stepped down from 250 to 206 ms: membrane voltage after the first (a) and the eighth (b) stimulus. (c) Membrane voltage at the probes P1–P4 (marked in a). 3D simulation: (d,e) Block of 100 × 100 × 40 cells with an ellipsoid of semi-axes 30, 30 and 17 cells, paced at the left face with the cycle length stepped down from 350 to 205 ms: cut-away views (front upper quarter removed) after the first (d) and the ninth (e) stimulus. (f) Membrane voltage at the probes P1–P4 (marked in d). Scale bars (b, e), 50 cells, i.e. 5 mm for a cell length of 100 µm. Dashed lines mark the stimuli, labeled with the beat number and the cycle length in ms; the shaded windows correspond to (a), (b), (d) and (e).

## 3 Discussion

Remodeling of ID nanostructure alters conduction, but its effect on conduction block in tissues could not be studied, because models that resolve this structure have been too costly for tissues beyond a cable [37, 38]. The hybrid model removes this limit by replacing only the junctional subsystem that dominates the cost with a learned flow map. Because the map predicts the two cleft currents, which collectively represent the GJ, ephaptic, capacitive, and ionic contributions of the ID, coupling between cells is preserved as a whole rather than reduced to a conductance. The hybrid model reproduced the FEM model across coupling regimes, ID geometries, pacing rates in cables, was validated in small 2D sheets, and in large tissues ran 10^4^-10^5^ times faster than the FEM model and within an order of magnitude of a monodomain model that omits the ID. Thus, resolving the ID no longer dominates the cost of a tissue simulation.

These are the first large-scale tissue simulations in which every junction resolves ID nanostructure. The hybrid model reproduces the experimentally observed dependence of conduction on perinexal width: widening speeds conduction with strong GJ coupling and slows conduction with reduced GJ coupling [13–15]. In heterogeneous tissues with reduced GJ coupling, the hybrid model shows that a region of widened clefts, as reported in AF [11], heart failure and ischemic heart disease [10], and after inflammatory vascular leak [12], creates a substrate for conduction block and reentry. Propagation through this cleft-expanded region is slowed and remains refractory when the next premature beat arrives, blocking conduction. Notably, GJ remodeling and cleft expansion occur together in these pathologies, and our simulations indicate that their combination (Figure 6), rather than either alone (Figure S5), sets the pro-arrhythmic substrate.

This dependence has consequences for therapy. Agents that enhance GJ coupling, block Na^+^ channels or stabilize perinexal adhesion [15] would be expected to act differently in tissue where ephaptic coupling carries a larger share of conduction, and the hybrid model makes these interactions testable in tissue for the first time. These predictions can be tested directly, for example by optical mapping during osmotic perinexal widening in hearts with reduced connexin-43 expression, and the recent direct demonstration of ephaptic coupling [18] makes such tests timely. In patient-specific models and digital twins that represent intercellular coupling by an averaged conductivity [50–52], the hybrid approach could capture arrhythmogenic regions driven by ID remodeling.

Importantly, the approach here is general. Because each map learns a single junction, its training cost is fixed by the junction, not the tissue, and the same pipeline yields maps for any ID geometry, including structures reconstructed from electron microscopy of a patient or disease model. More broadly, any multiscale system in which mechanistic units interact through an expensive, partially observed coupling element can be simulated in the same way: learn the coupling element, retain the mechanistic units, and assemble tissue of any size.

While this work establishes a junction-level surrogate that brings ID nanostructure into large-scale cardiac tissue simulations, we acknowledge key limitations in our study. First, each surrogate inherits any limitations of the FEM cable and associated training data, in which all ID-localized membranes are on the two cell ends [38]. This geometry is an approximation that does not hold in higher dimensions (Section 4.6), as the cleft currents depend nonlinearly on the geometry and channel density of a disk. Second, myocardial IDs are not arranged on a Cartesian lattice: branched myocytes bear several IDs of different size and structure, lateral IDs may mediate transverse conduction [39, 53], and in disease gap junctions lateralize and the perinexus widens [11]. Third, while hybrid tissue simulations were validated in a small 2D sheet with the FEM model, the 3D tissue predictions remain to be tested against experiments.

However, the surrogate itself is not tied to these simplifications. A single network conditioned on a parameter vector (i.e., ID geometry, GJ coupling and the share of a disk per connection), trained on single junctions, short cables and tissue-like voltage-clamp commands (Section 4.4), would define ID parameters for every junction, and cell-based models with realistic connectivity, such as the Kirchhoff network model [54] could naturally be deployed with this framework. Long term, this approach could bring ID nanostructure into organ-scale models, from in silico drug trials [55] to patient-specific virtual hearts that guide ablation [56] and digital twins [50–52], much like multiscale models that link calcium nanodomains to the whole heart [57].

## 4 Methods

### 4.1 Description of the reference FEM model

To link realistic ID nanostructure with tissue conduction, we previously developed a pipeline in which three-dimensional FEM meshes of the ID and the intercellular cleft were generated from transmission electron microscopy measurements, which were subsequently reduced to an equivalent electrical network of cleft compartments and embedded in a cable model of cardiac tissue [37]. This model was later extended [38, 58] to represent the spatial distribution of GJs and the nonuniform localization of ion channels and pumps on the ID membranes, together with electrodiffusion in the cleft, and used to simulate conduction for a wide range of parametrically defined ID geometries. This analysis identified GJ coupling, cleft geometry, and nanostructural heterogeneity as the dominant determinants of the cleft potential, the synchronization of the Na^+^ current and conduction velocity (CV), and showed that the plicate and interplicate intermembrane separations can either speed or slow conduction, depending on the strength of GJ coupling [38]. This FEM model is one of the most detailed representations of ID structure and channel organization that has been coupled to a tissue model, but its detail is computationally costly. Every junction is represented by hundreds of coupled cleft and ID potentials, cleft concentrations, and hundreds of active membrane patches (with associated ionic currents), with dynamics occurring on sub-millisecond timescales, such that numerical integration requires a suffciently small time step and a stiff integration scheme. As such, the dynamics of the junctions, rather than the cell axial membrane, dominate the computational cost of a simulation (described further below). This model has therefore so far been limited to one-dimensional cables of a few tens of cells, whereas a two-dimensional tissue large enough to sustain reentry contains 10^5^–10^6^ junctions [59].

Throughout this study, we use the most recently updated version of the FEM model [38] as the reference. The model resolves the detailed nanostructure of the ID (Figure 1a). The geometry of each ID is taken from a three-dimensional FEM mesh of the intercellular cleft (Figure 1b), parameterized by the ID diameter *d*, the interplicate and plicate intermembrane distances *d*_*ip*_ and *d*_*p*_, the plicate fold amplitude *A*_*f*_, and the interplicate height *h*_*ip*_ [38]. As detailed in our prior work [37], each mesh is reduced to an equivalent electrical network of *M*_disk_ = 100 compartments, in which a pre-junctional and a post-junctional ID membrane patch face a common cleft node (Figure 1b,d). The network also includes the spatial distribution of gap junctions, the nonuniform distribution of Na^+^ channels, inward-rectifier K^+^ channels, and the Na^+^*/*K^+^ ATPase over the ID membrane, and electrodiffusion of Na^+^, K^+^, Ca^2+^ and a generic anion A^−^ within the cleft. These ID networks are embedded in a one-dimensional cable of ventricular myocytes, in which each cell is represented by an axial membrane node whose ionic currents follow the O’Hara–Rudy model of the human ventricular myocyte (ORd11) [60].

We consider six ID geometries, meshes 1–6 (Figure 1c), which share *d* = 5 µm, *h*_*ip*_ = 3.2 µm and *A*_*f*_ = 0 nm and differ in the intermembrane distances *d*_*p*_ and *d*_*ip*_. Each mesh is combined with two values of the total GJ conductance *g*_gap_ of a junction, referred to as strong and reduced coupling.

#### Cell equations

We consider a cable of *N*_cell_ cells numbered *i* = 1, …, *N*_cell_ from left to right. Each cell has an axial membrane and two ID membranes, its left and right disks. Junction 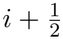 consists of the facing ID membranes of cells *i* and *i* + 1 and the cleft between them. Its equations are given in Supplemental Material. All interior junctions are discretized into *M*_disk_ ID membrane patches on both the pre- and post-junctional sides of the cleft, while the post-junctional (left) ID of cell 1 and the pre-junctional (right) ID of cell *N*_cell_ face the bulk with a single ID membrane.

Let 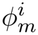 denote the axial potential of cell *i* and 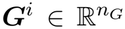 the ORd11 state variables of the axial membrane. Let 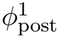 and 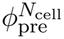 denote the potentials of the terminal disks. The governing equations for the interior cells *i* = 2, …, *N*_cell_ − 1 are

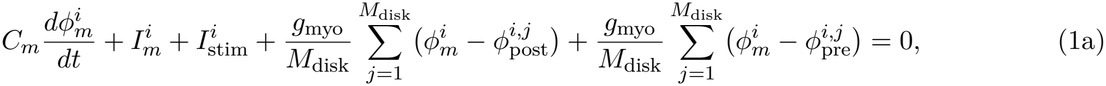

for the terminal cells,

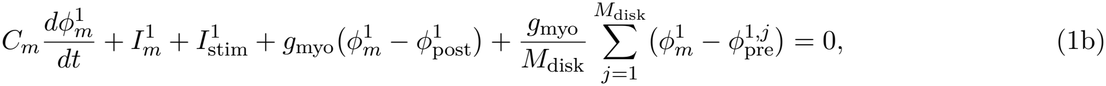

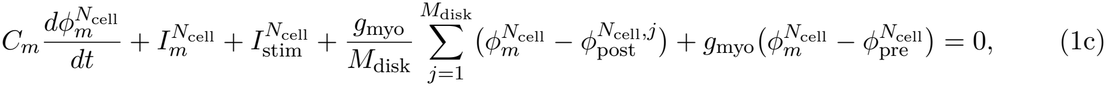

for the terminal disks,

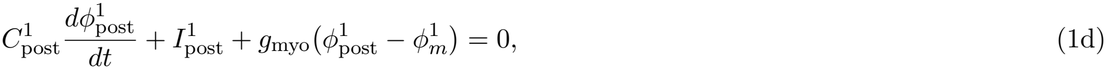

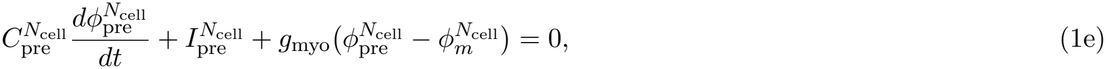

and for all cells *i* = 1, …, *N*_cell_,

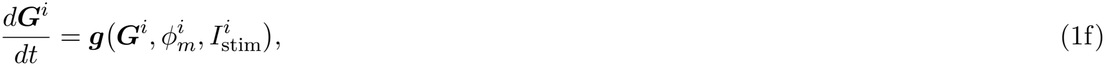

where *C*_*m*_ is the capacitance of the axial membrane, 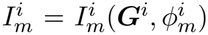 its ionic current and 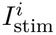 the applied stimulus. In the sums, 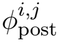 and 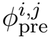 are the intracellular potentials of the *j*th ID membrane patches on the left and right IDs of cell *i*. These disks are the post-junctional side of junction 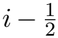 and the pre-junctional side of junction 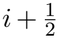. Each ID is connected to the axial node by the myoplasmic conductance *g*_myo_ (conductance of one-half the intracelllar space), such that each junctional ID membrane is connected to the axial node with conductance *g*_myo_*/M*_disk_. The terminal disks have the capacitances 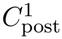 and 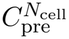 and the ionic currents 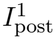 and 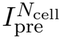 given in Section S7. The ORd11 state ***G***^*i*^ has *n*_*G*_ = 40 components: the gating variables, the intracellular ionic concentrations, and the state of the Ca^2+^/calmodulin-dependent protein kinase II (CaMK). The function 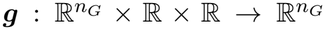 is the right-hand side of ORd11 model dynamics [60] with fixed extracellular concentrations and given stimulus current (carried by K^+^). Since the bulk extracellular space is grounded, 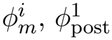 and 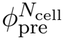 also represent transmembrane potentials.

#### Numerical solver

We solve Equation (1) and the junction equations by backward Euler for the circuit, with the ionic currents taken from the previous step. The potentials of all nodes then follow from one sparse linear system per step, whose constant matrix is factorized once per step size. The ORd11 gating variables are advanced by the Rush–Larsen method [61]. The other ionic states and the cleft concentrations are advanced by forward Euler with the potentials held fixed.

Importantly, almost all of the computational cost for a simulation of the FEM model arises from junction dynamics, for two reasons. First, the small cleft volumes result in cleft concentrations with stiff dynamics, requiring sufficiently small time steps. We advance them with 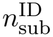 substeps per step. Second, a junction has about 10^4^ unknowns and a cell only *n*_*G*_ + 1 = 41, such that computational cost thus grows in proportion to the number of junctions.

Tissue-scale studies of arrhythmia require far larger domains. For example, recent two-dimensional atrial simulations use 512 × 512 to 1024 × 1024 cells (up to 12.8 × 12.8 cm, comparable to the area of both atria) [59]. With nearest-neighbor coupling, an *N*_*x*_ *× N*_*x*_ tissue contains 2*N*_*x*_(*N*_*x*_ *−*1) junctions, about 5.2 × 10^5^ for *N*_*x*_ = 512 and 2.1 × 10^6^ for *N*_*x*_ = 1024, corresponding to ~3.7 × 10^8^ and ~1.5 × 10^9^ junction potentials and cleft concentrations. This computational load makes it infeasible to account for the ID nanostructure in large tissue-scale simulations.

### 4.2 Observable dynamics of the junction

In Equation (1a), cell *i* interacts with each of its two junctions only through one sum, the current from its axial node into that junction. For junction 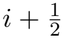, the currents from cells *i* and *i* +1 are the two cleft currents (purple arrows in Figure 1d)

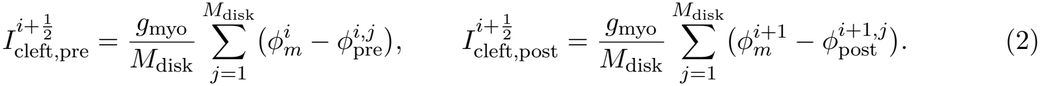

The two sums in Equation (1a) are thus 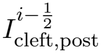 and 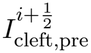, and the cell equations are closed once these currents are known. We therefore treat each junction as a dynamical system with a full state, an input and observables.

Let 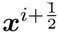 denote the full state of junction 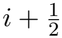,

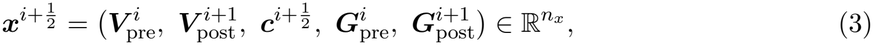

where 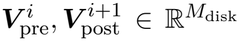 are the transmembrane potentials of the ID patches, 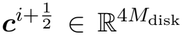 the concentrations of the four species in the cleft compartments, and 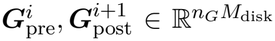 the ORd11 states of the ID patches. Here 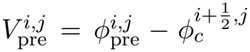 and 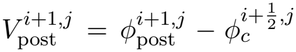, with 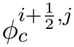 the potential of cleft compartment *j*. The full state thus has *n*_*x*_ = (2*n*_*G*_ + 6)*M*_disk_ = 86*M*_disk_ components. The ID and cleft potentials 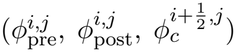 are not part of the full state, but rather follow from the full state (specifically the ID transmembrane potentials, 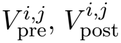 and the axial potentials 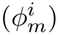 through a linear current balance at the ID and cleft nodes.

The input of junction 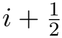 is the pair of axial potentials 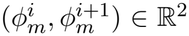, and its observables are the two cleft currents 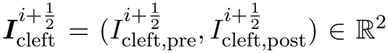. The junction equations of Section S7 then take the form

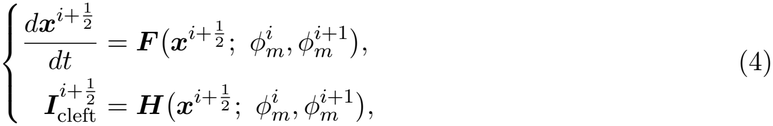

where 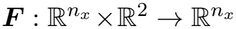 contains the currents of the ID membrane patches and the gap junctions, and the electrodiffusion in the cleft [38]. The output map 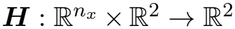 is Equation (2) with the ID potentials expressed through the full state and the input.

The observables alone do not determine their own evolution. By Equation (4), the observables at *t*_*n*+1_ depend on the full state at *t*_*n*_ and on the input over [*t*_*n*_, *t*_*n*+1_]. The cells provide the input but not the full state. Two observables cannot fix a full state of *n*_*x*_ = 8600 components. As different full states can give the same observables and evolve differently, the history of the observables provides the missing information. The longer the history, the less ambiguity it leaves about the full state. Under this condition, called *diminishing ambiguity* [44], a finite history of the observables and the input determines the next observables up to a small error.

The history must cover the slowest process that affects the observables. In this system, this process is the recovery of Na^+^ channels of the ID membrane patches from inactivation, which takes tens to hundreds of milliseconds [60]. A memory of this length is impractical. The ID membrane patches, however, follow nearly the same voltage history as its axial membrane. The Na^+^ availability of the axial membrane is (*hj*)^*i*^ = (0.99 *h*_*f*_ + 0.01 *h*_*s*_) *j*, where *h*_*f*_, *h*_*s*_ and *j* are ORd11 gating variables in ***G***^*i*^, which reasonably approximates Na^+^ channel availability of the ID patches. We therefore extend the input to the source

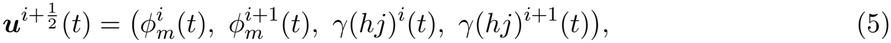

where *γ* = 100 is a numerical scaling factor. Unlike the axial potentials, (*hj*)^*i*^ does not enter Equation (4). It enters the observable dynamics only as information about the full state.

Let 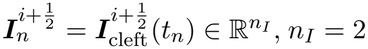 and 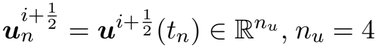, denote the observables and the source at time *t*_*n*_. We drop the superscript for a single junction. With a memory of *n*_*M*_ steps, the observables evolve by

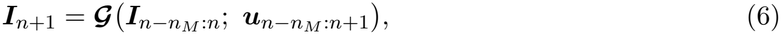

where 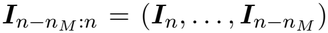 and 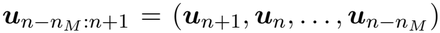 are the histories and 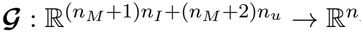is a finite-memory approximation of the observable dynamics. The source enters up to *t*_*n*+1_ because ***H*** depends on the axial potentials at *t*_*n*+1_. With *n*_*I*_ = 2 and *n*_*u*_ = 4, we have 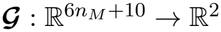.

### 4.3 Flow map learning model of the junction observables

The map ***G*** in Eqn. (6) is unknown, and therefore we learn ***G*** from data. The learned model must meet three requirements. First, it advances the observables from their history and that of the source, without the full state. Second, it describes a single junction. One model then serves all junctions of a tissue of any size. Third, it is computationally inexpensive enough to be evaluated at every junction in every step.

Neural networks have been used in several ways to model or solve dynamical systems. One class represents the right-hand side of an ODE by a network, as in neural ordinary differential equations (neural ODEs) [62]. This approach has been extended to learn the unknown components of differential–algebraic equations [63]. A second class embeds known equations in the training loss to solve forward and inverse problems, as in physics-informed neural networks [64, 65]. Neural operators, such as DeepONet [66] and the Fourier neural operator [67], learn maps between function spaces, for example, the solution operator of a parametric family of equations [68]. Recurrent networks and reservoir computers have been used to forecast chaotic systems [69, 70]. Autoencoders have been used to find coordinates in which the dynamics are approximately linear [71]. For partially observed systems, networks with memory learn the missing terms in the equations of the resolved variables [72–74]. For the junction, these methods face three difficulties. First, neural ODEs learn a continuous right-hand side. They inherit the stiffness of the cleft dynamics, which makes training difficult [75]. A trained model also depends on the integrator used in training [76]. Second, physics-informed networks and neural operators represent the solution on a whole domain. A model trained on a cable is then tied to the cable geometry. Third, networks with memory add missing terms to known equations of the resolved variables. No such equations exist for the cleft currents.

In this work, we use flow map learning (FML) [40]. Rather than learning a right-hand side or a solution on a domain, FML learns the evolution over a fixed time step, called the flow map, directly from trajectory data. It thus works from discrete data to a discrete model. FML was first introduced for autonomous systems [40] and later extended to parameterized systems [77], non-autonomous systems [78], partially observed systems [41], partial differential equations [79, 80] and stochastic systems [81, 82]. A detailed review of FML can be found in [83]. A memory-based FML model of one junction advances the observables from their history and that of the source, does not require equations of the observables, and its step is not limited by the stiffness of cleft dynamics. FML therefore meets all three requirements.

#### Flow map learning

For an unknown autonomous system *d* ***ξ*** */dt* = ***f*** (***ξ***), ***ξ*** ℝ^*d*^, where the right-hand side ***f*** : ℝ^*d*^ ℝ^*d*^ is not known, FML seeks to learn its flow map **Ψ** : ℝ^*d*^ ℝ^*d*^ instead of ***f***. The flow map governs the evolution of the solution over one step, that is, ***ξ*** (*t* _*n*+1_) = **Ψ**(***ξ*** (*t*_*n*_) with *t*_*n*+1_ = *t*_*n*_ + Δ*t*. When trajectory data are available, we split them into pairs ***ξ***^(*j*)^(*t*_*n*_), ***ξ***^(*j*)^(*t*_*n*+1_) separated by one step. Since the system is autonomous, the starting time *t*_*n*_ may diFFer between pairs. We then seek a function ***N*** : ℝ^*d*^ *→* ℝ^*d*^ that minimizes the loss

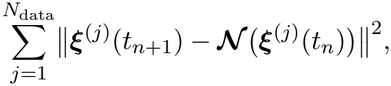

where *N*_data_ is the number of data pairs. Upon minimizing this loss, we obtain *N*≈**Ψ** and the predictive FML model

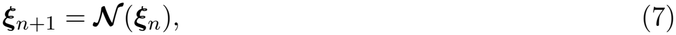

which predicts the solution once an initial state ***ξ***_0_ is given.

A notable extension of FML is the modeling of partially observed systems [41]. Let ***ξ*** = (***z, w***), where ***z*** ℝ^*m*^ are the observables and ***w ∈*** ℝ^*dℝm*^ the missing variables. When trajectory data are available only on ***z***, the work of [41] established that the dynamics of ***z*** can be modeled without any information on ***w***. Motivated by the Mori–Zwanzig formulation [84, 85], a discrete approximate Mori–Zwanzig formulation showed that a discrete model of ***z*** exists for *n ≥ n*_*M*_ in the form

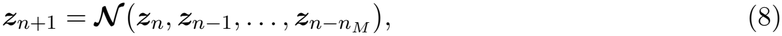

where *n*_*M*_ *≥* 0 is the number of memory steps and : 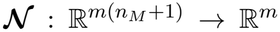 the flow map with memory. The case *n*_*M*_ = 0 corresponds to the fully observed system (7). For linear systems with linear observables, the work of [44] established that the observables satisfy a closed equation. The time derivative of the observables is a linear combination of their values at a finite number of delays. For nonlinear systems, a sufficiently long history determines the observables approximately under diminishing ambiguity, the condition behind Eqn. (6). For non-autonomous systems, the external inputs and their history enter the flow map as additional arguments [78]. This form has modeled the drag and lift on a cylinder under active flow control [43].

#### FML model

We adopt a neural network ***N***_*θ*_ to express the map ***G*** in Eqn. (6). Following [43, 78], the FML model of a junction takes the form

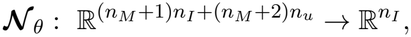

and evolves the observables according to

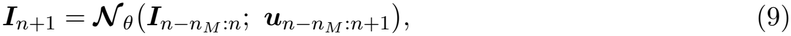

where *θ* denotes the parameters of the network. The model needs neither the full state nor the junction maps ***F*** and ***H***. The ID mesh properties and GJ coupling do not appear explicitly in the model. Each combination of the six meshes and the two couplings therefore gives its own trained model, with its own parameters *θ*. All of them share the same network structure and the same memory.

The most important parameter of the model is the memory length *T*_*M*_ = *n*_*M*_ Δ *t*. The model uses the step Δ *t* = 0.02 ms, ten times the step of the FEM model. For this step, numerical tests with increasing memory length, reported in Section 2.1, show that *n*_*M*_ = 50, that is, *T*_*M*_ = 1 ms, gives the smallest error among the memories tested. The same tests support the inclusion of *γ* (*hj*) in the source. With *n*_*M*_ = 50, we have ***N***_*θ*_ : ℝ^310^ → ℝ^2^. The training of ***N***_*θ*_ is described in Section 4.4.

### 4.4 Construction of the FML model

#### Data generation

To construct the FML model in Eqn. (9), training data are generated by the FEM model with the step Δ *t*_FEM_ = 0.002 ms and 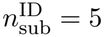. Simulations of the FEM model for different pacing rate conditions also provide the validation data (described below). The training data must cover the conditions that the model responds to in tissue simulations. For each combination of ID mesh and GJ coupling, the procedure consists of the following steps:

1. Run cables of 50 and 10 cells with the same mesh at every junction (Figure 7a). Cell 1 is paced, and the voltages of all cells evolve freely. In the 50-cell cable, most junctions respond to a fully developed propagating wave. The 10-cell cable adds the upstrokes shaped by the stimulus and by the far end of the cable, as every junction of this cable lies within a few cells of the cable end.
2. Run voltage-clamp (VC) cases of a single junction, with its two cells clamped to the commands 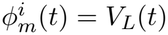 and 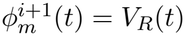 (Figure 7b). The VC cases cover voltage pairs distinct from the cable trajectories. Such pairs arise in tissue simulation and capture local perturbations of the cell voltages.
3. Record the cleft currents and the source of every junction every Δ *t*. The Na^+^ availability (*hj*) is reconstructed from each voltage trace by integrating the ORd11 gates *h*_*f*_, *h*_*s*_ and *j* with the Rush–Larsen scheme on this grid. Current and voltage quantities are kept in physical units, mV and µ A, respectively.

**Figure 7:**
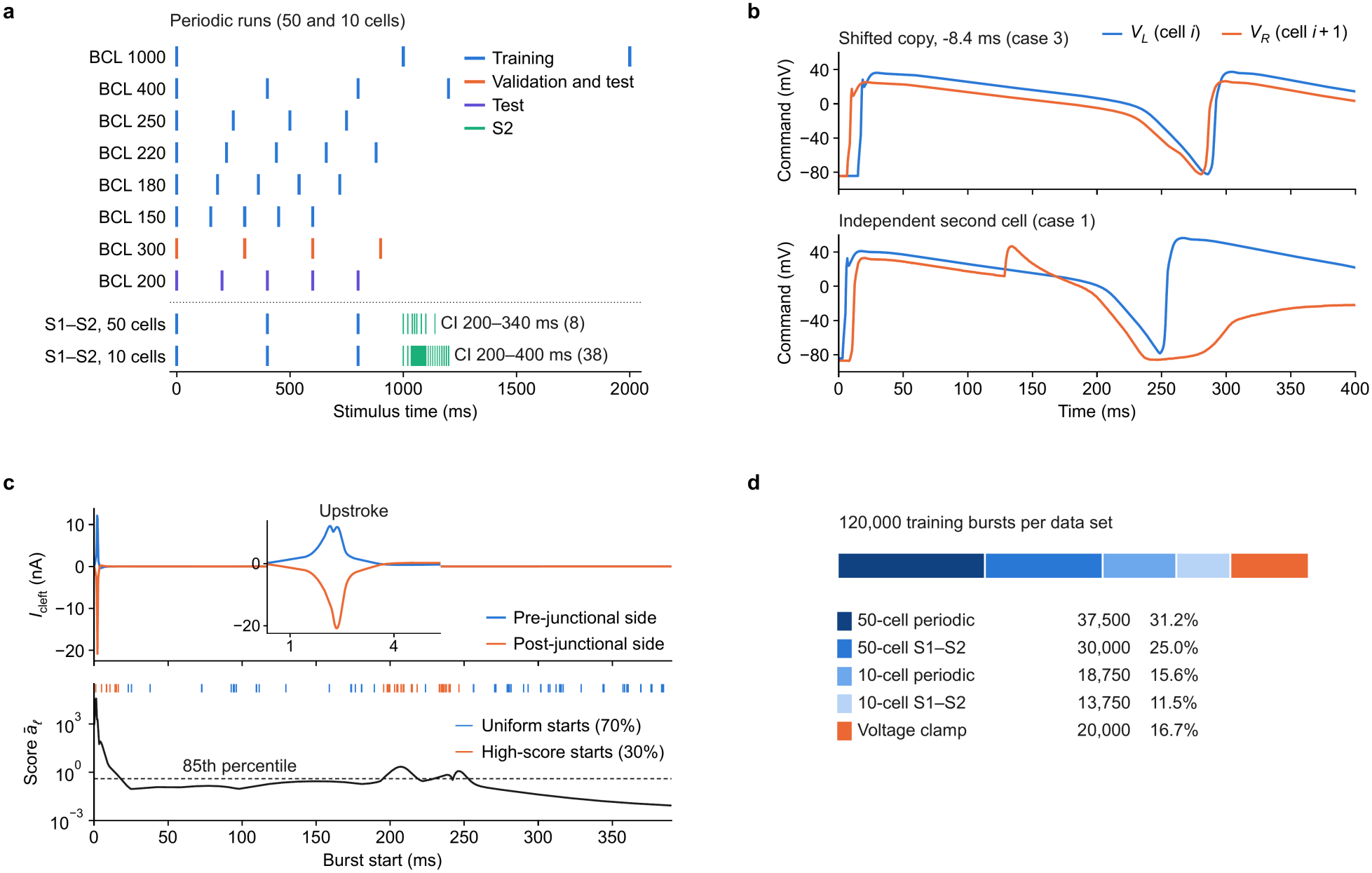
Overview of the data generation process. (a) Stimulus times of the FEM cable runs, colored by their role. The periodic runs are made with 50 and 10 cells. The S1–S2 runs vary the CI of the S2 beat. (b) Clamp commands of two VC cases for mesh 1 and strong coupling. Top: the second command is a shifted and perturbed copy of the first. Bottom: the second command is independent. (c) Burst sampling at one junction of a 10-cell training run, for mesh 3, strong coupling, and the first beat at BCL 400 ms. Top: cleft currents of the two sides of the junction, with the upstroke enlarged. Bottom: score *ā*_*ℓ*_ of each burst start and its 85th percentile. The ticks mark the drawn starts. (d) Composition of the 120 000 training bursts of each data set.

The procedure yields a time history of ***I***_*n*_ and ***u***_*n*_ for every junction of every run.

Both cables are paced periodically at six basic cycle lengths (BCLs), 1000, 400, 250, 220, 180 and 150 ms. They are also paced with an S1–S2 protocol, in which three S1 beats at BCL 400 ms are followed by a premature S2 beat. The coupling interval (CI) of the S2 takes eight values between 200 and 340 ms in the 50-cell cable. In the 10-cell cable, it takes 38 values between 200 and 400 ms, spaced by 2 ms between 236 and 276 ms, close to the threshold for refractoriness. Periodic runs at two further BCLs, 300 and 200 ms, are held out for validation and testing.

A VC case runs for 400 ms. The command voltage for one cell starts with a randomized FEM action potential of the same mesh and coupling. One or two premature events follow in the plateau, late repolarization, early diastole, or the recovered state. The command voltage for the second cell is usually a shifted and perturbed copy of the first, which gives small voltage differences of either sign. In other cases, the second cell command voltage is independent. Each data set has 300 VC cases. Of these, 235 are used for training, and the other 65 are held out.

The FEM model is needed only for this offline data generation process. VC runs also offer a way to train models for realistic tissue at the cost of a single junction, with commands that mimic the local conditions of two- and three-dimensional tissue.

#### Training data

The memory *n*_*M*_ = 50 and *n*_*R*_ = 5 recurrent steps require training data of length *n*_*L*_ = *n*_*M*_ + 1 + *n*_*R*_ = 56. We therefore sample bursts of *n*_*L*_ consecutive samples of one junction from the trajectory data. The first *n*_*M*_ +1 samples of a burst serve as the initial condition, and the last *n*_*R*_ are predicted in training. With Δ *t* = 0.02 ms, a burst spans 1.12 ms, of which the initial condition covers 1.02 ms and the predicted part 0.1 ms.

Each data set contains 120 000 training bursts. Of these, 67 500 come from the 50-cell runs, 32 500 from the 10-cell runs and 20 000 from the VC cases (Figure 7d). The VC fraction is kept small, such that the physiological cable trajectories dominate. In the cable runs, the bursts are spread evenly over the junctions. In the S1–S2 runs, they are restricted to [*t*_S2_ – 60, *t*_S2_ + 340] ms around the premature beat. From each VC training case, we cut 100 bursts. Half of them lie within [*t*_*e*_ –1, *t*_*e*_ + 40] ms of a randomly chosen protocol event *t*_*e*_. The 20 000 VC bursts are drawn from this pool.

Within a junction of a cable run, the starts of the bursts are not drawn uniformly. The upstroke occupies only a few milliseconds of each cycle but dominates the cleft currents, such uniform sampling would rarely contain the upstroke. We therefore score every sample *n* of a junction by its activity

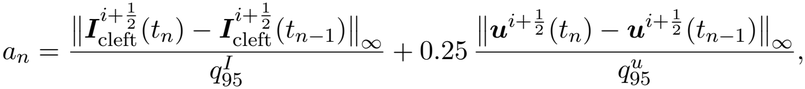

where 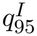 and 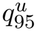 are the 95th percentiles of the two increments of that junction over the run. A burst starting at sample *l* receives the score 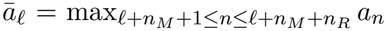, the largest activity among its predicted samples. For each junction, 30 % of the sample starts are drawn from the starts with *ā*_*l*_ above the 85th percentile, and the remain 70 % are sampled uniformly (Figure 7c).

### Training

We adopt a fully connected feedforward network for ***N*** _*θ*_. The input layer contains the nodes of 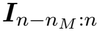 and 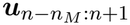, resulting in 310 nodes. The output layer contains the two nodes of ***I***_*n*+1_. In between, we employ four hidden layers of 80 nodes each, with rectified linear unit (ReLU) activations. The network thus has layer widths {310, 80, 80, 80, 80, 2}. Neither the input nor the output is normalized. In contrast with ohmic currents, the two cleft current outputs are not constrained to be equal and opposite, as current also flows through the ID membranes and from the cleft to the bulk.

To train the network, we use the first *n*_*M*_ + 1 samples of each burst as the initial condition, ***Î***_*v*_ = ***I***_*v*_ for *v* = *l*, …, *l* + *n*_*M*_, where *l* is the first sample of the burst. We then march the FML model forward *n*_*R*_ times. After each step, the predicted current replaces the observed one in the history, while the source is taken from the data. That is, for *r* = 1,.., *n*_*R*_,

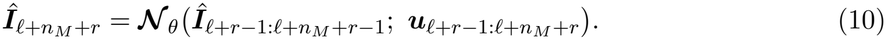

We then compute the mean squared error between the predictions ***Î*** and the data ***I***, averaged over the *n*_*R*_ steps and over the set *B* of training bursts,

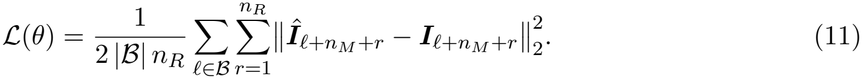

This is the recurrent loss that we minimize to train the network. The gradient is propagated through the fed-back predictions. The network is thus trained on the inputs it receives in simulations, where its own outputs fill the current history.

The network is trained with the Adam optimizer [86] to minimize the recurrent loss (Eqn. 11) over *n*_*R*_ = 5 recurrent steps. Training is conducted for 60 000 epochs with batch size 4096, in float64 arithmetic. An exponential learning rate schedule is used, with initial rate 5 × 10^−4^ and decay factor 0.9999 per epoch. The gradient norm is clipped at 5 × 10^−4^. If the training loss stays above 20 times its best value for 10 consecutive epochs, the best parameters are restored, the learning rate is halved, and the optimizer state is reset. We keep the parameters with the lowest loss on 50 validation bursts from the held-out 50-cell run at BCL 300 ms, computed with a longer rollout of *n*_*R*_ + 10 = 15 steps.

Following, we train *K* = 5 networks 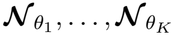 for every combination of ID mesh and GJ coupling, from different random initializations, and use them as an ensemble. The mean of the ensemble gives the next currents,

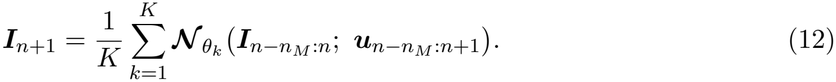

All members receive the same input, and the mean is appended to the shared history of the currents. In Section 4.5, ***N***_*θ*_ denotes this mean.

### 4.5 Description of the hybrid model

In the hybrid model, each junction is replaced by the FML model (Eqn. 9) with the parameters learned for the corresponding ID mesh and GJ coupling. The cell axial membranes retain their respective equations (Eqn. 1). Let 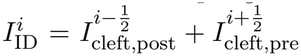 denote the current that cell draws from its two junctions, with the cleft currents given by the FML model. At the terminal cells, *I*_ID_ also contains the current into the terminal disk, 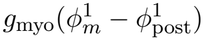 or 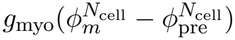.

Let 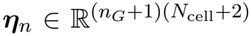 denote the cell unknowns, 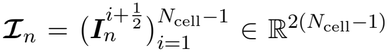 the cleft currents of all junctions and 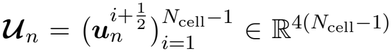 the sources of all junctions at time *t*_*n*_ = *n* Δ *t*. The cell unknowns ***ε***_*n*_ are the potential and the ORd11 state of every axial membrane and of the two terminal disks. One step of the hybrid model solves the system

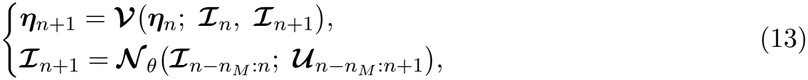

where 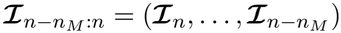 and 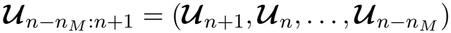 are the histories, and ***N***_*θ*_ acts on each junction separately, mapping 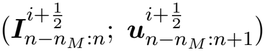 as in Eqn. (12). The newest source, ***U***_*n*+1_ = ***U***_*n*+1_(***η***_*n*+1_), is the source (Eqn. 5) evaluated with the axial potentials and the ORd11 gates of ***η***_*n*+1_. The map : 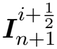 is the numerical solution of the cell equations, Eqn. (1), over one step, which allows for consideration of multiple integration schemes (described below). This integration is an operator splitting method that uses the cleft currents _*n*_ at *I*_*n*_ and *I*_*n*+1_ at *t*_*n*+1_, respectively, in the sums of Eqns. (1a)–(1c). The two equations in Eqn. 13 are coupled, as the FML model input includes the cell potentials at *t*_*n*+1_ through the source, and the cell potential includes the cleft currents.

#### Cell substeps

The map ***V*** uses *n*_sub_ substeps *t*_*n*,*k*_ = *t*_*n*_ + *k δ t, k* = 0, …, *n*_sub_, of length *δ t* = Δ*t*/*n*_sub_. On the substeps, the cleft currents are interpolated linearly

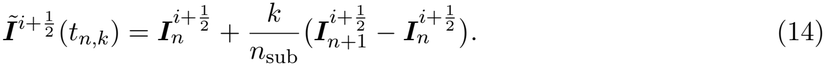

The axial potentials are advanced by the discretization of Eqns. (1a)–(1c),

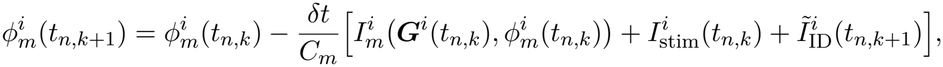

Where 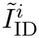 is the interpolated cleft currents. The ionic current is evaluated at the start of the substep and the cleft currents at its end. The terminal-disk equations, Eqns. (1d) and (1e), are discretized by backward Euler. Each terminal cell then results in a linear system for its axial and disk potentials. The ORd11 states, Eqn. (1f), are advanced with 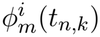, by the Rush–Larsen method for the gating variables and forward Euler for the other state variables. The histories of the FML model are updated only at the times *t*_*n*_.

#### Coupling schemes

We consider two coupling schemes, which di!er in the determination of *I*_*n*+1_ in *V*.

1. explicit performed a single iteration of a predictor-corrector scheme. The predictor defines 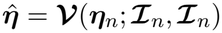, such that the FML model at 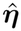 defines 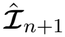, and the corrector is therefore 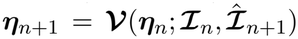. If the corrector changes any axial potential by more than 5 mV relative to the predictor, the predictor is kept.
2. Implicit solves Eqn. 13 iteratively using the implicit scheme described below.

After each step, the FML model is evaluated at ***η***_*n*+1_ to define *I*_*n*+1_, and the new currents and sources are appended to the histories. All runs begin at *t* = 0 with the cells at rest. The current histories are then zero, since the cleft currents vanish at rest, and the source histories take their resting values. The initial currents *I*_0_ are the outputs of the FML model for these histories. A run is stopped if any axial potential leaves [150, 300] mV.

We remark that the two schemes differ in cost and stability. A step of explicit costs two integrations of the cell equations and two evaluations of the FML model at all junctions, and implicit repeats both until Eqn. 13 is solved. For *n*_sub_ = 1, explicit is one fixed-point iteration of backward Euler and implicit is backward Euler. For a cleft current that is linear in the cell potential with slope *φ >* 0, explicit is therefore stable for *κ* Δ*t* < *C*_*m*_, whereas implicit is stable for any step.

#### Solution of the implicit scheme

To solve Equation (13) using the implicit scheme, we take the axial potentials 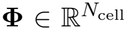 as the unknowns. The other cell states follow from ***V***. In these unknowns, the residual 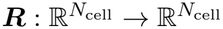 is

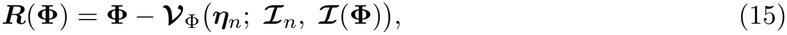

where ***V***_φ_ is the axial potential part of ***V*** and ***I***(**φ**) the outputs of the FML model with the target-time potentials set to **φ**. Each evaluation of ***R*** integrates the step again from the stored state ***η***_*n*_.

The residual is reduced by linearized steps. Let **φ**^*i*^ and *R*^*i*^ denote the entries of cell *i*. The step

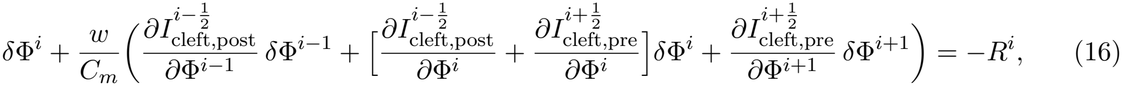

where 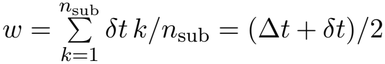 is the weight of the endpoint currents in the *n*_sub_ substeps.

At the terminal cells, the terms of the missing junction are dropped. The derivatives of the cleft currents are taken through the network, and those of the ionic currents are neglected. Each junction contributes the derivatives of its two outputs with respect to the potentials of its two cells. The system is therefore tridiagonal in the cable. In two- and three-dimensional tissues, each row couples a cell to its neighbors. The system is solved by LU factorization for up to 100 cells and otherwise by the generalized minimal residual method (GMRES) [88].

Each linearized step is followed by a backtracking line search, which halves the step up to ten times until ‖***R***‖_2_ decreases. If no step is accepted, the potentials are instead updated by the relaxed fixed-point step **φ** − *β****R***. The factor *β* starts at 0.5 and is halved whenever the residual decreases by less than 5%. We call a trial one evaluation of ***R*** together with the derivatives of the network. The iteration starts from the EXPLICIT predictor and stops when ‖***R***‖_∞_ < 1 × 10^−3^ mV.

### 4.6 Numerical experiments

We test the hybrid model at increasing scale. In cables, the FEM model provides a reference, and we compare the two models for varying pacing rates and in heterogeneous cables with mixed ID geometries. We also perform validation simulations in a small two-dimensional sheet. However, in large two- and three-dimensional tissues, simulation of the FEM model is infeasible. We simulate the hybrid model in two- and three-dimensional tissues with regional heterogeneity in ID geometries. Finally, we compare the computational cost of the hybrid model with that of the FEM model and of a monodomain model.

#### Cable simulations

##### Cable simulations

All cable simulations compare the hybrid model with the FEM model [38] on the same ID mesh and GJ coupling. Unless stated otherwise, the cable has 50 cells and is homogeneous with the same ID mesh at every junction. The cable is paced at cell 1 for ten beats. We perform validation simulations at the two held-out BCLs, 300 and 200 ms. The hybrid model uses the IMPLICIT scheme with *n*_sub_ = 4, that is, *δ t* = 0.005 ms, and the five-member ensemble.

We assess agreement between the hybrid surrogate model and FEM reference in the axial potentials, the cleft currents, and conduction velocity (CV). For the potential, we measure the root-mean-square error (RMSE) of 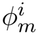 over all 50 cells and the whole run, in mV. The squared error is averaged over all cells and all time samples before the square root is taken, so the RMSE is normalized per cell and does not grow with the number of cells. The same measure, over all cells, is used in every simulation in which the hybrid model is compared with the FEM model. For the cleft currents, we measure the RMSE of 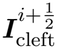 over both sides of the junctions 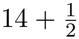 and 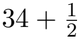 and over the whole run, normalized by the peak 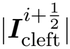 of the reference. CV is measured on the last beat, as the distance of 3 mm between cells 11 and 41 divided by the difference of their activation times. The activation time of cell *i* is the first upward crossing of 0 mV by 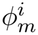, linearly interpolated between samples.

CV restitution measures the hybrid model over the whole range of pacing rates. For each combination of ID mesh and GJ coupling, we perform a decreasing pacing period protocol, in which the cable is paced for five beats at each BCL. The BCL decreases from 1000 ms to 300 ms in steps of 100 ms, and then in steps of 10 ms until conduction no longer follows every stimulus (i.e., the loss of 1:1 capture). At each BCL, CV of the last beat is compared with that of the FEM reference. Here, the FEM reference uses the adaptive time step of, with Δ*t*_FEM_ = 0.002 ms during the 50 ms after each stimulus, Δ*t*_FEM_ = 0.02 ms for the rest of the cycle and 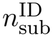 from 5 to 14.

Simulations in the heterogeneous cable assess a more physiological setting, in a cable with junctions with different ID geometries. The mesh of each of its 49 junctions is drawn independently and uniformly from the six meshes, and the junctions of the corresponding hybrid model use the parameters learned for its own mesh. The same mesh spatial distribution is used for the FEM reference and the hybrid model, for both GJ couplings and both BCLs. We compare the local CV for the two models: For *i* = 2, …, *N*_cell_ − 1, the local CV measurement follows from the activation times 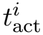,

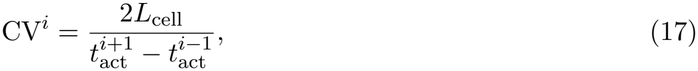

where *L*_cell_ = 100 µm is the cell length. The relative error of the hybrid model is 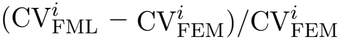.

#### Tissue simulations

The tissue simulations use a 100 × 100 sheet and a 100 × 100 × 40 block of identical cells. Simulation of a smaller 20 × 20 sheet is performed in both the FEM and hybrid model for validation, described in Section 4.7. In the higher-dimensional tissues, each cell is joined to its four or six neighbors by a junction, such that the sheet has 19 800 junctions and the block 1 182 000. To account for differences between the one-dimensional cable, in which cells have only two connections, in the sheet, each junction carries the current of half a disk. In the block, the two junctions along the pacing direction carry half the current of a disk each and the four lateral junctions a quarter each. The ID membrane currents and the GJ current of a junction are scaled by its share. In the cell equations, 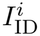 then sums the cleft currents of the four or six junctions of cell *i*, each scaled in the same way. The outer faces are sealed.

In the tissues with regional heterogeneity, the tissues contain a central region of widened clefts. The tissue has reduced GJ coupling throughout. Junctions whose two cells lie in the central region have properties of mesh 4, with *d*_*p*_ = *d*_*ip*_ = 50 nm, and all others mesh 1, with *d*_*p*_ = *d*_*ip*_ = 10 nm. The region is a circle of radius 30 cells in the sheet and an ellipsoid with semi-axes of 30, 30 and 17 cells in the block. Each tissue is paced ten times at its left edge or face, with a stimulus of 50 μA μF^−1^ for 2 ms. A decremental pacing protocol is applied: The sheet receives four beats at 250, two at 220, and four at 206 ms. The block receives one beat each at 350, 300, 270, 250, 230 and 210 ms, then four at 205 ms. The potential is recorded at four probes on the line through the center of the region along the pacing direction (Figure 6a,d). They lie before the region (P1), in its proximal half (P2), at its center (P3) and beyond it (P4). The tissue simulations use the EXPLICIT scheme with *n*_sub_ = 1, that is, *δ t* = Δ*t* = 0.02 ms, and one member of each ensemble instead of the mean, run on a GPU with single precision.

Noting that the networks were trained with the wave traveling from cell *i* to cell *i*+1, to account for the asymmetry of directional propagation, in the higher dimensional tissues, the “roles” of the two cells are therefore assigned at the onset of each activation, based on when the first crosses −80 mV. The cell with the higher potential then takes the role of cell *i*, and the histories of the junction are swapped when the roles change.

#### Computational cost

The cost of the hybrid model is compared with that of the FEM model and of the standard monodomain model (without clefts). To benchmark the monodomain model, every junction is replaced by an ohmic gap junction with the conductance of the reduced coupling, 73.5 nS, and each cell is a single ORd11 compartment with the capacitance of the whole cell. All models run on one workstation with a 16-core CPU, 31 GB of memory and a consumer GPU, with full specifications are given in Table S2. The FEM model runs in its MATLAB implementation with Δ*t*_FEM_ = 0.001 ms. The hybrid and monodomain models run in JAX on the CPU and the GPU with Δ*t* = 0.02 ms, with both IMPLICIT and EXPLICIT. For the 3-D tissues, they also run in a separate CUDA implementation of the same schemes. The timed cases are the 50-cell cable, with mesh 1, reduced GJ coupling, BCL 400 ms, the five-member ensemble and *n*_sub_ = 4, and the two tissues.

For the FEM model, the sheet, with about 6 × 10^6^ nodes, fits into the memory of the workstation, with a peak of 22 GB, and is run in full. However, the block would require about 1.5 TB, such that FEM run time is therefore estimated by scaling the time of a 20⨯ 20⨯ 8 block of the same shape, with 8880 junctions, by the number of junctions. The cost per junction and simulated millisecond, 314 ms, 331 ms and 327 ms in the cable, the sheet and the small block, supports this scaling.

We time the second simulated millisecond of the first beat, excluding the setup, the LU factorization of the FEM model and the compilation of the JAX code. The CUDA implementation compiles its kernels before the loop and is timed over the first two milliseconds. The timed interval lies in the stimulus, where implicit needs 3.0–3.3 trials per step, against 2.3–2.6 on average over whole beats. The times of implicit are thus upper bounds. The cost per step of the FEM model does not depend on the state, and that of the hybrid model depends on it only through the number of trials. The times therefore extrapolate linearly to longer runs.

### 4.7 Declaration of use of AI in study and manuscript writing

Generative AI tools were used to assist with debugging and refactoring the code to improve its readability, including removing redundant code and reorganizing it into modules, and to check grammar and improve the clarity of the manuscript. The authors reviewed and verified all AI-assisted changes and take responsibility for the final code and manuscript.

## Acknowledgments

This project was supported by funding from the National Institutes of Health R01HL165751 (SHW).

## Author contributions

**RS and XL:** These authors contributed equally to this work. Conceptualization, Methodology, Writing – original draft, Visualization, Validation, Software, Methodology, Investigation. **NM:** Conceptualization, Methodology, Writing - review & editing. **DX:** Conceptualization, Methodology, Resources, Supervision, Writing - review & editing. **SHW:** Conceptualization, Funding acquisition, Methodology, Project administration, Resources, Supervision, Writing - original draft.

