## Supplemental Material for "A hybrid biophysical–neural network model links nanoscale cardiac junctions to tissue conduction"

### Supplementary Material

#### Notation

Table S1 lists the symbols used in the paper. Spatial indices are superscripts, namely the cell $i$, the junction $i+\frac{1}{2}$ and the compartment $j$. The time level $n$ is a subscript. At junction $i+\frac{1}{2}$, the pre-junctional side belongs to cell $i$ and the post-junctional side to cell $i+1$. Thus pre is the right disk of a cell and post its left disk. Bold symbols are vectors and vector-valued maps.

Table S1: Notation.

| Symbol | Meaning | Unit |
|---|---|---|
| **Indices and counts** | | |
| $i = 1, \ldots, N_{\text{cell}}$ | cell of the cable | – |
| $i+\frac{1}{2}$, $i-\frac{1}{2}$ | junction (ID and cleft) between cells $i$ and $i+1$, and between $i-1$ and $i$ | – |
| $j = 1, \ldots, M_{\text{disk}}$; $j_1, j_2, j_3$ | compartment of an ID: a cleft node and the pre- and post-junctional ID patches facing it; several compartments | – |
| $M_{\text{disk}} = 100$ | compartments per ID | – |
| $s \in \{\text{Na}^+, \text{K}^+, \text{Ca}^{2+}, \text{A}^-\}$; $z_s$ | ionic species in the cleft, $\text{A}^-$ a generic anion; valence | – |
| $N_x$ | cells per side of a square sheet (cost estimate) | – |
| $n$, $t_n = n\,\Delta t$ | time level and its time | –, ms |
| $k = 0, \ldots, n_{\text{sub}}$; $t_{n,k} = t_n + k\,\delta t$ | substep index and substep times; $t_{n,0} = t_n$, $t_{n,n_{\text{sub}}} = t_{n+1}$ | –, ms |
| $\ell$ | first sample of a training burst | – |
| $\nu$, $r$ | sample index and rollout step of the recurrent training (Equation (10)) | – |
| **Potentials** | | |
| $\phi_m^i$ | axial potential of cell $i$, also its transmembrane potential; $\phi_m$ labels the source that holds the axial potentials only | mV |
| $\phi_{\text{pre}}^{i,j}$, $\phi_{\text{post}}^{i,j}$ | intracellular potential of the $j$th pre-junctional (right disk) and post-junctional (left disk) ID compartment of cell $i$ | mV |
| $\phi_c^{i+\frac{1}{2},j}$ | potential of cleft compartment $j$ of junction $i+\frac{1}{2}$ | mV |
| $\phi_{\text{post}}^1$, $\phi_{\text{pre}}^{N_{\text{cell}}}$ | potentials of the two terminal disks, which face the bulk | mV |
| $\boldsymbol{\Phi}$ | axial potentials of all cells at $t_{n+1}$, the unknowns of the implicit scheme | mV |
| $V_L(t)$, $V_R(t)$ | voltage-clamp commands of $\phi_m^i$ and $\phi_m^{i+1}$ | mV |
| **Currents and circuit** | | |
| $C_m$ | capacitance of the axial membrane | $\mu$F |
| $C_{\text{post}}^1$, $C_{\text{pre}}^{N_{\text{cell}}}$ | capacitances of the terminal disks | $\mu$F |
| $g_{\text{myo}}$ | myoplasmic conductance between the axial node and one ID; $g_{\text{myo}}/M_{\text{disk}}$ to each compartment | mS |
| $I_m^i$ | ionic current of the axial membrane of cell $i$ | $\mu$A |
| $I_{\text{post}}^1$, $I_{\text{pre}}^{N_{\text{cell}}}$ | ionic currents of the terminal disks | $\mu$A |
| $I_{\text{stim}}^i$ | stimulus current, carried by $\text{K}^+$ | $\mu$A |

*continued on the next page*

*Table S1, continued*

| Symbol | Meaning | Unit |
|---|---|---|
| $\boldsymbol{I}_{\text{cleft}}^{i+\frac{1}{2}} = (I_{\text{cleft,pre}}^{i+\frac{1}{2}}, I_{\text{cleft,post}}^{i+\frac{1}{2}})$ | cleft currents of junction $i+\frac{1}{2}$: from cell $i$ into the pre-junctional and from cell $i+1$ into the post-junctional side (Equation (2)) | $\mu$A |
| $I_{\text{ID}}^{i}$; $\tilde{I}_{\text{ID}}^{i}$ | current drawn by cell $i$ from its junctions (Section 3.5); the same with the interpolated currents | $\mu$A |
| $\Delta t_{\text{FEM}}$; $n_{\text{sub}}^{\text{ID}}$ | time step of the FEM ID model; cleft-concentration substeps per step: the smallest number for a stable explicit update with a safety factor of four, and at least five | ms; – |
| **Junction state and ionic model** | | |
| $\boldsymbol{x}^{i+\frac{1}{2}}$ | full state of junction $i+\frac{1}{2}$ (Equation (3)), in $\mathbb{R}^{n_x}$, $n_x = (2n_G+6)M_{\text{disk}} = 86M_{\text{disk}}$ | – |
| $\boldsymbol{F}$, $\boldsymbol{H}$ | right-hand side and output map of the junction system (Equation (4)) | – |
| $V_{\text{pre}}^{i,j}$, $V_{\text{post}}^{i+1,j}$; $\boldsymbol{V}_{\text{pre}}^{i}$, $\boldsymbol{V}_{\text{post}}^{i+1}$ | transmembrane potentials of the ID patches of compartment $j$ of junction $i+\frac{1}{2}$, $\phi_{\text{pre}}^{i,j} - \phi_{c}^{i+\frac{1}{2},j}$ and $\phi_{\text{post}}^{i+1,j} - \phi_{c}^{i+\frac{1}{2},j}$; the same over $j$ | mV |
| $\boldsymbol{c}^{i+\frac{1}{2}}$; $c_s^j$, $\boldsymbol{c}^j$ | cleft concentrations of junction $i+\frac{1}{2}$, in $\mathbb{R}^{4M_{\text{disk}}}$; concentration of species $s$ in compartment $j$, and those of compartment $j$ | mM |
| $\boldsymbol{G}^{i}$ | ORd11 state of the axial membrane of cell $i$ (gates, intracellular concentrations, CaMK), in $\mathbb{R}^{n_G}$, $n_G = 40$ | – |
| $\boldsymbol{G}_{\text{pre}}^{i,j}$, $\boldsymbol{G}_{\text{post}}^{i,j}$ | ORd11 states of the ID patches of compartment $j$ | – |
| $\boldsymbol{g}$ | right-hand side of the ORd11 state equations | – |
| **Junction equations (Section S7)** | | |
| $A^j$, $C^j$, $\text{vol}^j$ | area and capacitance of each ID patch of compartment $j$; cleft volume of compartment $j$ | $\mu\text{m}^2$, $\mu$F, $\mu\text{m}^3$ |
| $g_c^{jj'}$, $g_b^j$ | conductance between cleft nodes $j$ and $j'$; between cleft node $j$ and the bulk | mS |
| $g_{\text{gap}}$, $\sigma_{\text{gap}}^j$ | total gap-junctional conductance of a junction; its weight in compartment $j$ | mS, – |
| $g_{\text{ID}}^{jj'}$, $\rho_{\text{ID,cleft}}$ | conductance between ID patches $j$ and $j'$ of the same side; its ratio to $g_c^{jj'}$ $(=1)$ | mS, – |
| $I_X$; $\omega_X$, $\sigma_X^j$ | whole-cell ORd11 current $X$; its weight on the two disks and its weight in compartment $j$ | $\mu$A; – |
| $R_{\text{cg}}$, $C_{\text{tot}}$ | ratio of capacitive to geometric membrane area in ORd11 $(=2)$; capacitance of the geometric area of a cell | –, $\mu$F |
| $I_{\text{pre}}^{i,j}$, $I_{\text{post}}^{i+1,j}$; $I_{\text{pre},s}^{i,j}$, $I_{\text{post},s}^{i+1,j}$ | ionic currents of the ID patches of compartment $j$; their parts carried by species $s$ | $\mu$A |
| $J_s^j$ | electrodiffusive current of species $s$ leaving cleft node $j$ | $\mu$A |
| $E_s^{jj'}$, $E_s^{j\text{b}}$ | Nernst potentials of species $s$ between cleft nodes $j$ and $j'$ and between node $j$ and the bulk | mV |
| $c_{s,\text{b}}$, $\boldsymbol{c}_{\text{b}}$ | bulk concentrations | mM |
| $R$, $T$, $F$ | gas constant, temperature, Faraday constant | – |
| $h_f$, $h_s$, $j$ | fast and slow inactivation gates and recovery gating variables of $I_{\text{Na}}$ in ORd11 ($j$ here is a gating variable) | – |
| $(hj) = (0.99\,h_f + 0.01\,h_s)\,j$ | $\text{Na}^+$ channel availability | – |

*continued on the next page*

*Table S1, continued*

| Symbol | Meaning | Unit |
|---|---|---|
| $\gamma = 100$ | scaling of $(hj)$ in the source | – |
| **Flow map** | | |
| $\boldsymbol{\xi} = (\boldsymbol{z}, \boldsymbol{w})$, $\boldsymbol{f}$ | state of a dynamical system, its observables and missing variables; right-hand side | – |
| $\boldsymbol{\Psi}$ | flow map of an autonomous system over one step (Section 3.3) | – |
| $N_{\text{data}}$ | number of data pairs in the review of FML (Section 3.3) | – |
| $\boldsymbol{\mathcal{N}}$ | function that approximates a flow map (Equation (7)); with memory, Equation (8) | – |
| $\boldsymbol{\mathcal{G}}$ | map of the observable dynamics of a junction (Equation (6)) | – |
| $\boldsymbol{\mathcal{N}}_\theta$, $\theta$ | network of the FML model of a junction and its parameters; in the hybrid model, the mean of the ensemble | – |
| $K = 5$; $\theta_k$ | ensemble size; parameters of member $k$ (Equation (12)) | – |
| $n_I = 2$, $n_u = 4$ | dimensions of the cleft currents and of the source of a junction (Equation (9)) | – |
| $\boldsymbol{u}^{i+\frac{1}{2}}$ | source of junction $i + \frac{1}{2}$, $(\phi_m^i, \phi_m^{i+1}, \gamma(hj)^i, \gamma(hj)^{i+1})$ (Equation (5)) | – |
| $\boldsymbol{I}_n$, $\boldsymbol{u}_n$ | $\boldsymbol{I}_n^{i+\frac{1}{2}} = \boldsymbol{I}_{\text{cleft}}^{i+\frac{1}{2}}(t_n)$ and $\boldsymbol{u}_n^{i+\frac{1}{2}} = \boldsymbol{u}^{i+\frac{1}{2}}(t_n)$; superscript dropped for a single junction (Sections 3.2 to 3.4) and kept in the hybrid model (Section 3.5) | $\mu$A, – |
| $\boldsymbol{I}_{n-n_M:n}$, $\boldsymbol{u}_{n-n_M:n+1}$ | histories $(\boldsymbol{I}_n, \ldots, \boldsymbol{I}_{n-n_M})$ and $(\boldsymbol{u}_{n+1}, \ldots, \boldsymbol{u}_{n-n_M})$ | – |
| $\Delta t$ | step of the FML model (0.02 ms) | ms |
| $n_M$, $T_M = n_M \Delta t$ | memory in steps and its length | –, ms |
| $n_R$ | recurrent steps of the training loss | – |
| $n_L = n_M + 1 + n_R$ | length of a training burst, in samples | – |
| **Data and training** | | |
| $a_n$ | activity of sample $n$ | – |
| $q_{95}^I$, $q_{95}^u$ | 95th percentiles of the increments of the cleft currents and of the source | – |
| $\bar{a}_\ell$ | score of the burst that starts at $\ell$ | – |
| $t_{\text{S2}}$, $t_e$ | onset of the premature S2 stimulus; a protocol event of a voltage-clamp case | ms |
| $\hat{\boldsymbol{I}}_\nu$ | rolled-out prediction (Equation (10)) | $\mu$A |
| $\mathcal{L}(\theta)$, $\mathcal{B}$ | recurrent loss (Equation (11)); set of training bursts | – |
| **Hybrid model** | | |
| $n_{\text{sub}}$, $\delta t = \Delta t / n_{\text{sub}}$ | substeps per step and their length | –, ms |
| $\boldsymbol{\eta}_n$ | cell unknowns at $t_n$: potentials and ORd11 states of all cells and of the two terminal disks, in $\mathbb{R}^{(n_G+1)(N_{\text{cell}}+2)}$ in the cable | – |
| $\boldsymbol{\mathcal{I}}_n$ | cleft currents of all junctions at $t_n$, in $\mathbb{R}^{2(N_{\text{cell}}-1)}$ in the cable | $\mu$A |
| $\boldsymbol{\mathcal{U}}_n$ | sources of all junctions at $t_n$, in $\mathbb{R}^{4(N_{\text{cell}}-1)}$ in the cable (Equation (13)) | – |
| $\tilde{\boldsymbol{I}}^{i+\frac{1}{2}}(t_{n,k})$ | linear interpolation of the cleft currents on the substeps (Equation (14)) | $\mu$A |
| $\boldsymbol{\mathcal{V}}$, $\boldsymbol{\mathcal{V}}_\Phi$ | numerical solution of the cell equations over one step, $\mathbb{R}^{(n_G+1)(N_{\text{cell}}+2)} \times \mathbb{R}^{2(N_{\text{cell}}-1)} \times \mathbb{R}^{2(N_{\text{cell}}-1)} \to \mathbb{R}^{(n_G+1)(N_{\text{cell}}+2)}$ (Equation (13)); its axial-potential part, with values in $\mathbb{R}^{N_{\text{cell}}}$ | – |

*continued on the next page*

*Table S1, continued*

| Symbol | Meaning | Unit |
|---|---|---|
| $\hat{\boldsymbol{\eta}}$, $\hat{\boldsymbol{\mathcal{I}}}_{n+1}$ | predicted state of EXPLICIT and the cleft currents at it | – |
| EXPLICIT, IMPLICIT | coupling schemes (Section 3.5) | – |
| $\boldsymbol{R}(\boldsymbol{\Phi})$; $R^i$, $\Phi^i$, $\delta\Phi^i$ | residual of IMPLICIT, $\mathbb{R}^{N_{\text{cell}}} \to \mathbb{R}^{N_{\text{cell}}}$ (Equation (15)); its entry, the unknown potential and the linearized step of cell $i$ (Equation (16)) | mV |
| $w = (\Delta t + \delta t)/2$ | weight of the currents at $t_{n+1}$ in the linearized step (Equation (16)) | ms |
| $\beta$ | relaxation factor of the fixed-point fallback of IMPLICIT | – |
| $\kappa$ | slope of a cleft current that depends linearly on the cell potential | mS |
| **ID geometry** | | |
| $d$ | ID diameter | $\mu$m |
| $d_{ip}$, $d_p$ | interplicate and plicate intermembrane distances | nm |
| $A_f$ | fold amplitude | nm |
| $h_{ip}$ | interplicate height | $\mu$m |
| **Conduction** | | |
| $t^i_{\text{act}}$ | activation time of cell $i$: first upward crossing of 0 mV | ms |
| CV, $\text{CV}^i$ | conduction velocity; local conduction velocity of cell $i$ (Equation (17)) | cm/s |
| $L_{\text{cell}}$ | cell length (100 µm) | $\mu$m |
| BCL, CI | basic cycle length; S1–S2 coupling interval | ms |

#### Junction equations in the FEM reference model

The equations of junction $i + \frac{1}{2}$ below define $\boldsymbol{F}$ and $\boldsymbol{H}$ in Equation (4). They follow Eq. S1 of the FEM model [38], which builds on [37], as implemented in the code that generated the training data. In this section, the cleft quantities carry only the compartment index $j, j' = 1, \ldots, M_{\text{disk}}$. The ionic species are $s \in \{\text{Na}^+, \text{K}^+, \text{Ca}^{2+}, \text{A}^-\}$, with valence $z_s$.

**Parameters.** Compartment $j$ has a pre- and a post-junctional ID patch and a cleft volume $\text{vol}^j$. Each patch has the area $A^j$ and a capacitance $C^j$ proportional to $A^j$, as for the axial membrane. Neighboring cleft nodes are coupled through the conductances $g_c^{jj'} = g_c^{j'j}$, with $g_c^{jj'} = 0$ for non-neighbors. Cleft node $j$ is coupled to the grounded bulk through $g_b^j$. Both conductances are computed from the FEM mesh and the extracellular resistivity. The gap junctions have the total conductance $g_{\text{gap}}$, distributed over the compartments with weights $\sigma^j_{\text{gap}}$, $\sum_j \sigma^j_{\text{gap}} = 1$. ID patches of the same side are coupled through the intracellular space of the disk with the conductances $g^{jj'}_{\text{ID}} = \rho_{\text{ID,cleft}}\, g_c^{jj'}$, $\rho_{\text{ID,cleft}} = 1$.

**Membrane currents.** Let $I_X(\boldsymbol{G}, V; \boldsymbol{c})$ be the whole-cell current $X$ of ORd11 for the state $\boldsymbol{G}$, the transmembrane potential $V$ and the extracellular concentrations $\boldsymbol{c}$. It is the ORd11 current density times $R_{\text{cg}} C_{\text{tot}}$, where $C_{\text{tot}}$ is the capacitance of the geometric membrane area of the cell and $R_{\text{cg}} = 2$ the ratio of capacitive to geometric area in ORd11. Each ORd11 current $X$ lies on the two disks with the weight $\omega_X$ and on the axial membrane with the weight $1 - \omega_X$. Here $\omega_{\text{Na}} = 0.7$ for the fast $\text{Na}^+$ current, $\omega_{\text{K1}} = \omega_{\text{CaL}} = \omega_{\text{NaK}} = 0.2$, $\omega_X$ is the share of the two disks in the membrane area of the cell for $I_{\text{to}}$, $I_{\text{Kr}}$ and $I_{\text{Ks}}$, and $\omega_X = 0$ for all other currents. On a disk, current $X$ is distributed over the compartments with weights $\sigma_X^j$, $\sum_j \sigma_X^j = 1$. The weights follow the nonuniform distributions of the $\mathrm{Na}^+$ channels, the inward-rectifier $\mathrm{K}^+$ channels and the $\mathrm{Na}^+/\mathrm{K}^+$ ATPase [38]. For all other currents, they are proportional to $A^j$. The ionic current of the pre-junctional patch of compartment $j$ is

$$I_{\mathrm{pre}}^{i,j} = \frac{1}{2} \sum_X \omega_X \, \sigma_X^j \, I_X\big(\boldsymbol{G}_{\mathrm{pre}}^{i,j}, V_{\mathrm{pre}}^{i,j}; \boldsymbol{c}^j\big),$$

where $\boldsymbol{c}^j$ holds the cleft concentrations of compartment $j$ and the factor $\frac{1}{2}$ divides the disk weight between the two disks of the cell. The current $I_{\mathrm{post}}^{i+1,j}$ is defined in the same way. We write $I_{\mathrm{pre},s}^{i,j}$ for the part of $I_{\mathrm{pre}}^{i,j}$ carried by species $s$, with $I_{\mathrm{pre,A^-}}^{i,j} = 0$ and $\sum_s I_{\mathrm{pre},s}^{i,j} = I_{\mathrm{pre}}^{i,j}$. The states of the patches obey

$$\frac{d\boldsymbol{G}_{\mathrm{pre}}^{i,j}}{dt} = \boldsymbol{g}\big(\boldsymbol{G}_{\mathrm{pre}}^{i,j}, V_{\mathrm{pre}}^{i,j}; \boldsymbol{c}^j\big), \qquad \frac{d\boldsymbol{G}_{\mathrm{post}}^{i+1,j}}{dt} = \boldsymbol{g}\big(\boldsymbol{G}_{\mathrm{post}}^{i+1,j}, V_{\mathrm{post}}^{i+1,j}; \boldsymbol{c}^j\big), \tag{S1}$$

where $\boldsymbol{g}$ is the ORd11 right-hand side of Equation (1f), evaluated with the currents of the patch, with the cleft concentrations $\boldsymbol{c}^j$ in place of the fixed extracellular ones and without stimulus. Every patch has its own intracellular concentrations, and the patches do not exchange intracellular ions. The axial membrane carries the remaining weights, $I_m^i = \sum_X (1-\omega_X)\, I_X\big(\boldsymbol{G}^i, \phi_m^i; \boldsymbol{c}_{\mathrm{b}}\big)$, with the bulk concentrations $\boldsymbol{c}_{\mathrm{b}}$. Each terminal disk in Equations (1d) and (1e) carries the share $\frac{1}{2}\omega_X$ of every current, with its own ORd11 states and the bulk concentrations. Its capacitance is proportional to the total area $\sum_j A^j$ of the ID mesh.

**Node equations.** Kirchhoff's current law at the pre-junctional, post-junctional and cleft nodes of compartment $j$ gives

$$C^j \frac{dV_{\mathrm{pre}}^{i,j}}{dt} + I_{\mathrm{pre}}^{i,j} + \frac{g_{\mathrm{myo}}}{M_{\mathrm{disk}}}\big(\phi_{\mathrm{pre}}^{i,j} - \phi_m^i\big) + g_{\mathrm{gap}}\sigma_{\mathrm{gap}}^j\big(\phi_{\mathrm{pre}}^{i,j} - \phi_{\mathrm{post}}^{i+1,j}\big) + \sum_{j'} g_{\mathrm{ID}}^{jj'}\big(\phi_{\mathrm{pre}}^{i,j} - \phi_{\mathrm{pre}}^{i,j'}\big) = 0, \tag{S2}$$

$$\begin{aligned} C^j \frac{dV_{\mathrm{post}}^{i+1,j}}{dt} &+ I_{\mathrm{post}}^{i+1,j} + \frac{g_{\mathrm{myo}}}{M_{\mathrm{disk}}}\big(\phi_{\mathrm{post}}^{i+1,j} - \phi_m^{i+1}\big) + g_{\mathrm{gap}}\sigma_{\mathrm{gap}}^j\big(\phi_{\mathrm{post}}^{i+1,j} - \phi_{\mathrm{pre}}^{i,j}\big) \\ &+ \sum_{j'} g_{\mathrm{ID}}^{jj'}\big(\phi_{\mathrm{post}}^{i+1,j} - \phi_{\mathrm{post}}^{i+1,j'}\big) = 0, \end{aligned} \tag{S3}$$

$$\sum_s J_s^j = C^j \frac{dV_{\mathrm{pre}}^{i,j}}{dt} + I_{\mathrm{pre}}^{i,j} + C^j \frac{dV_{\mathrm{post}}^{i+1,j}}{dt} + I_{\mathrm{post}}^{i+1,j}, \tag{S4}$$

with $V_{\mathrm{pre}}^{i,j} = \phi_{\mathrm{pre}}^{i,j} - \phi_c^{i+\frac{1}{2},j}$ and $V_{\mathrm{post}}^{i+1,j} = \phi_{\mathrm{post}}^{i+1,j} - \phi_c^{i+\frac{1}{2},j}$ (Equation (3)). Here $J_s^j$ is the electrodiffusive current of species $s$ that leaves the cleft node,

$$J_s^j = \frac{1}{4}\Big[\sum_{j'} g_c^{jj'}\big(\phi_c^j - \phi_c^{j'} - E_s^{jj'}\big) + g_b^j\big(\phi_c^j - E_s^{j\mathrm{b}}\big)\Big],$$

with the Nernst potentials $E_s^{jj'} = \frac{RT}{z_s F}\ln\big(c_s^{j'}/c_s^j\big)$ and $E_s^{j\mathrm{b}} = \frac{RT}{z_s F}\ln\big(c_{s,\mathrm{b}}/c_s^j\big)$, where $c_{s,\mathrm{b}}$ is the bulk concentration. As in the implementation of [38], each conductance is shared equally among the four species. The cleft concentrations obey

$$z_s F\, \mathrm{vol}^j \frac{dc_s^j}{dt} = I_{\mathrm{pre},s}^{i,j} + I_{\mathrm{post},s}^{i+1,j} - J_s^j. \tag{S5}$$

**Reduction to Equation (4).** The cleft node has no capacitance of its own, so Equations (S2) to (S4) are differential–algebraic. Adding them eliminates the capacitive and ionic currents, and the gap-junctional terms cancel:

$$\sum_s J_s^j + \frac{g_{\text{myo}}}{M_{\text{disk}}}(\phi_{\text{pre}}^{i,j} - \phi_m^i + \phi_{\text{post}}^{i+1,j} - \phi_m^{i+1}) + \sum_{j'} g_{\text{ID}}^{jj'}(\phi_{\text{pre}}^{i,j} - \phi_{\text{pre}}^{i,j'} + \phi_{\text{post}}^{i+1,j} - \phi_{\text{post}}^{i+1,j'}) = 0. \quad \text{(S6)}$$

With $\phi_{\text{pre}}^{i,j} = V_{\text{pre}}^{i,j} + \phi_c^{i+\frac{1}{2},j}$ and $\phi_{\text{post}}^{i+1,j} = V_{\text{post}}^{i+1,j} + \phi_c^{i+\frac{1}{2},j}$, Equation (S6) is a linear system for the cleft potentials. Its matrix is symmetric positive definite: $(1 + 2\rho_{\text{ID,cleft}})$ times the graph Laplacian of $g_c^{jj'}$ plus the diagonal $g_b^j + 2g_{\text{myo}}/M_{\text{disk}}$. Given $\boldsymbol{x}^{i+\frac{1}{2}}$, $\phi_m^i$ and $\phi_m^{i+1}$, the system therefore determines $\phi_c^j$, and with it $\phi_{\text{pre}}^{i,j}$ and $\phi_{\text{post}}^{i+1,j}$. Equations (S2) and (S3) then give $dV_{\text{pre}}^{i,j}/dt$ and $dV_{\text{post}}^{i+1,j}/dt$, Equation (S5) gives $dc_s^j/dt$ and Equation (S1) the patch states. Together they form $\boldsymbol{F}$. The output map $\boldsymbol{H}$ is Equation (2) evaluated with these potentials.

#### Cable simulations

Figures S1 and S2 show the closed-loop traces summarized in Figure 2. The surrogates are for mesh 1, with the memories $T_M = 0.24$, 0.5 and 1 ms. Each memory is combined with the full source $\boldsymbol{u}$ of Equation (5) and with a source that contains only the axial potentials, $\phi_m$. In the panels, $\boldsymbol{u}$ is labeled "V + h·j" and $\phi_m$ "V only". Figure S3 shows the membrane potentials and cleft currents of the 50-cell cable with heterogeneous ID geometry of Section 1.4.

#### Validation in a small two-dimensional sheet

We compared the surrogate with the FEM model in a sheet of $20 \times 20$ cells with four connections per cell (400 cells, 760 junctions), with mesh 1 ($d_p = d_{ip} = 10$ nm) at every junction, for strong and for reduced GJ coupling (Figure S4). In the FEM model, each cell has four ports: the ID-localized channels of a cell are divided equally among its four connections (one quarter of a disk per connection), each connection carries the full GJ conductance, and faces without a neighbor are closed by a lumped boundary-disk patch. The FEM model is integrated with the scheme of the training data ($\Delta t = 0.002$ ms, cleft substep 0.0004 ms) and saved every 0.02 ms. In the surrogate, each connection is represented by the flow map of the same mesh and coupling (full source $\boldsymbol{u}$, $T_M = 1$ ms, a single network), trained on cables in which each connection carries half a disk. To match the four-port convention, the predicted cleft currents entering the cell equations are scaled by 1/2 while the GJ current is kept in full. The unscaled outputs are stored in the history of the network, such that its inputs stay within the training range. Faces without a neighbor are closed by an active terminal-disk patch with the area of the mesh (30.7 µm$^2$). The surrogate is integrated with the implicit scheme of the 1-D simulations (IMPLICIT, $n_{\text{sub}} = 4$, coupling tolerance $1 \times 10^{-3}$ mV). In both models, the left column of cells is stimulated with 50 µA µF$^{-1}$ for 2 ms and one beat of 400 ms is simulated.

The surrogate reproduces the activation map and the action potentials of the reference (Figure S4). The conduction velocity is 40.2 against 38.3 cm s$^{-1}$ with strong GJ coupling (4.8 %) and 16.0 against 16.3 cm s$^{-1}$ with GJ reduced coupling (−1.7 %). The action potential duration, from the upward crossing of 0 mV to the downward crossing of −75 mV averaged over the 400 cells, is shorter in the surrogate by 8.3 and 17.2 ms. The RMSE of the potential over all 400 cells and the 400 ms run, normalized per cell as in the cable (Section 3.6), is 5.29 and 10.53 mV. The error is nearly uniform over the sheet, with a per-cell RMSE of 5.2 mV–5.4 mV with strong and 10.2 mV–10.8 mV with reduced GJ coupling. Activation agrees within about 0.1 ms, and about 95 %

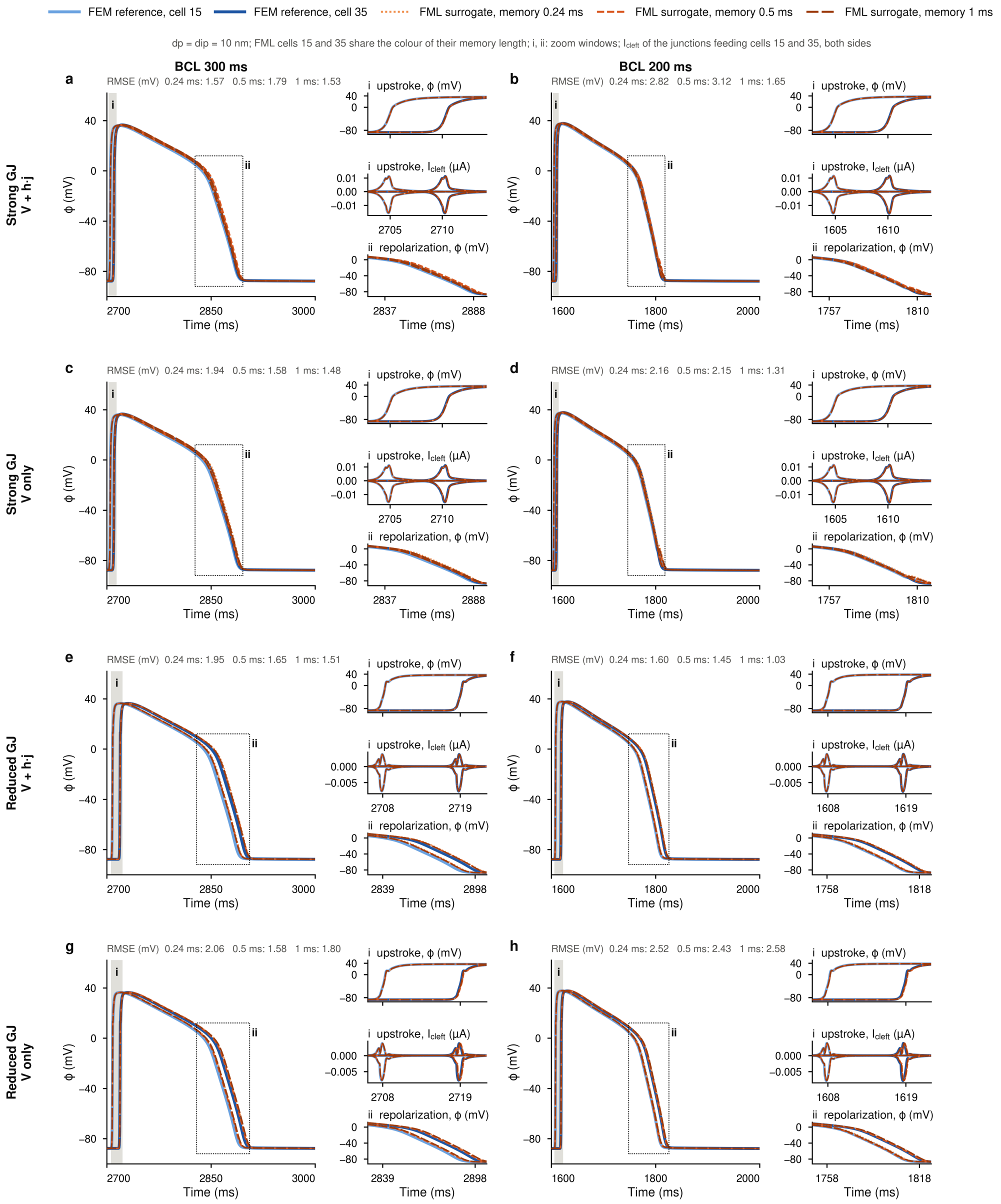


Figure S1: Effect of the memory on the closed-loop cable, mesh 1. Rows: strong and reduced gap-junctional coupling, each with the full source (V + h·j) and with the axial potentials only (V only). Columns: BCL 300 ms and 200 ms. Each panel shows the last beats of cells 15 and 35 for the FEM reference and for the surrogates with $T_M = 0.24$, 0.5 and 1 ms. The insets enlarge the upstroke (i), with the potential and the cleft currents of junctions $14 + \frac{1}{2}$ and $34 + \frac{1}{2}$, and repolarization (ii). The RMSE over all 50 cells is given above each panel.

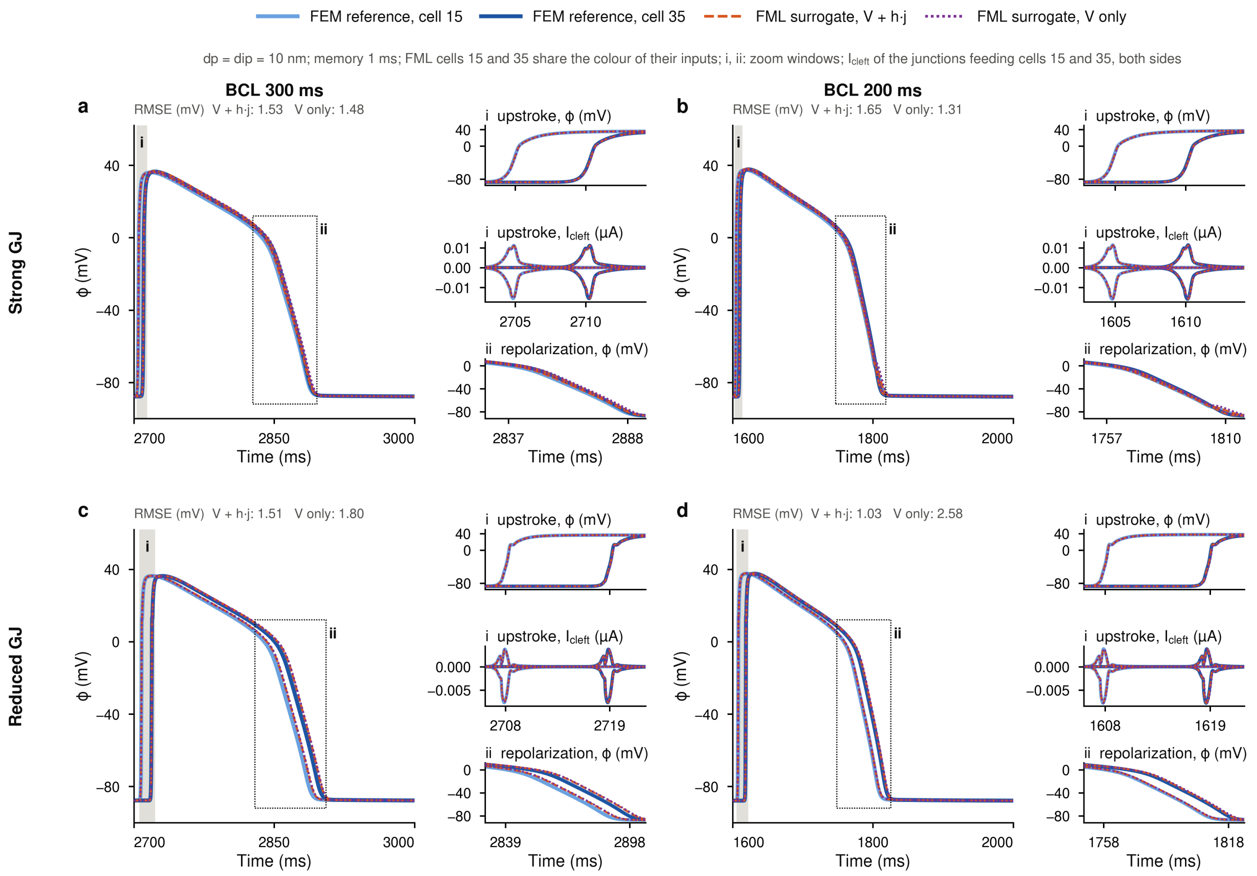


Figure S2: Effect of the source on the closed-loop cable, mesh 1, memory 1 ms. The surrogates use the full source (V + h·j) or the axial potentials only (V only). (a,b) Strong and (c,d) reduced gap-junctional coupling, at BCL 300 ms (a,c) and 200 ms (b,d). Layout as in Figure S1.

of the squared error occurs during repolarization. The larger RMSE than in the cable (1 mV–2 mV) therefore reflects the shorter action potential of the surrogate, likely due to the scaling of the cleft currents by 1/2 used to place the cable-trained maps in a sheet with four connections per cell.

#### Control simulations for the three-dimensional tissue block

Two control cases in comparison with the heterogeneous 3D tissue block with reduced GJ coupling of Figure 6d–f were performed. Simulations were run with the same lattice, pacing protocol, coupling scheme, and probes (Figure S5): (i) a homogeneous tissue (with mesh 1 at every junction) and reduced GJ coupling and (ii) the heterogeneous tissue but strong GJ coupling. In contrast with the heterogeneous 3D tissue block with reduced GJ coupling, in both control cases, every stimulus conducts as a plane wave, and there is no local conduction block observed.

#### Benchmark hardware

The wall times of Table 1 are measured on one local workstation, one run at a time and with nothing else running. Table S2 lists its hardware and software. The table gives a peak and a measured floating-point rate for the CPU and the GPU, in double (FP64) and single (FP32) precision. The peak rate is the product of the number of cores, the clock, and the floating-point operations per

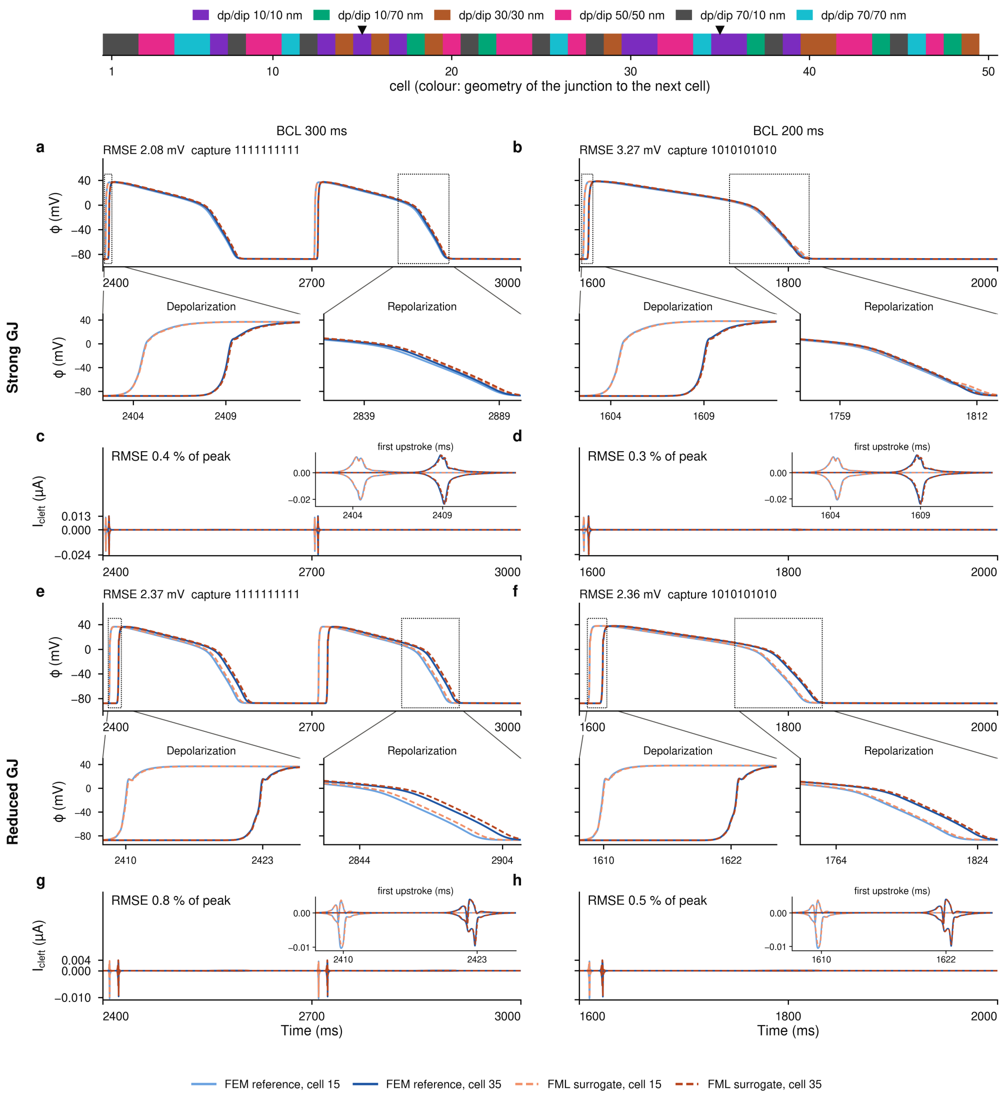


Figure S3: Closed-loop simulation of a 50-cell cable with heterogeneous ID geometry. Top: mesh of each junction along the cable. The color at cell $i$ is that of junction $i + \frac{1}{2}$, and the triangles mark cells 15 and 35. (a,b,e,f) Last beats of cells 15 and 35 for strong (a,b) and reduced (e,f) gap-junctional coupling, at BCL 300 ms (a,e) and 200 ms (b,f), with depolarization and repolarization enlarged. Above each panel are the RMSE over all 50 cells and the capture pattern of the ten beats, with 1 if the beat propagates along the cable and 0 if not. (c,d,g,h) Cleft currents of the junctions feeding cells 15 and 35 over the same window, with the first upstroke enlarged. The RMSE is given in percent of the peak current.

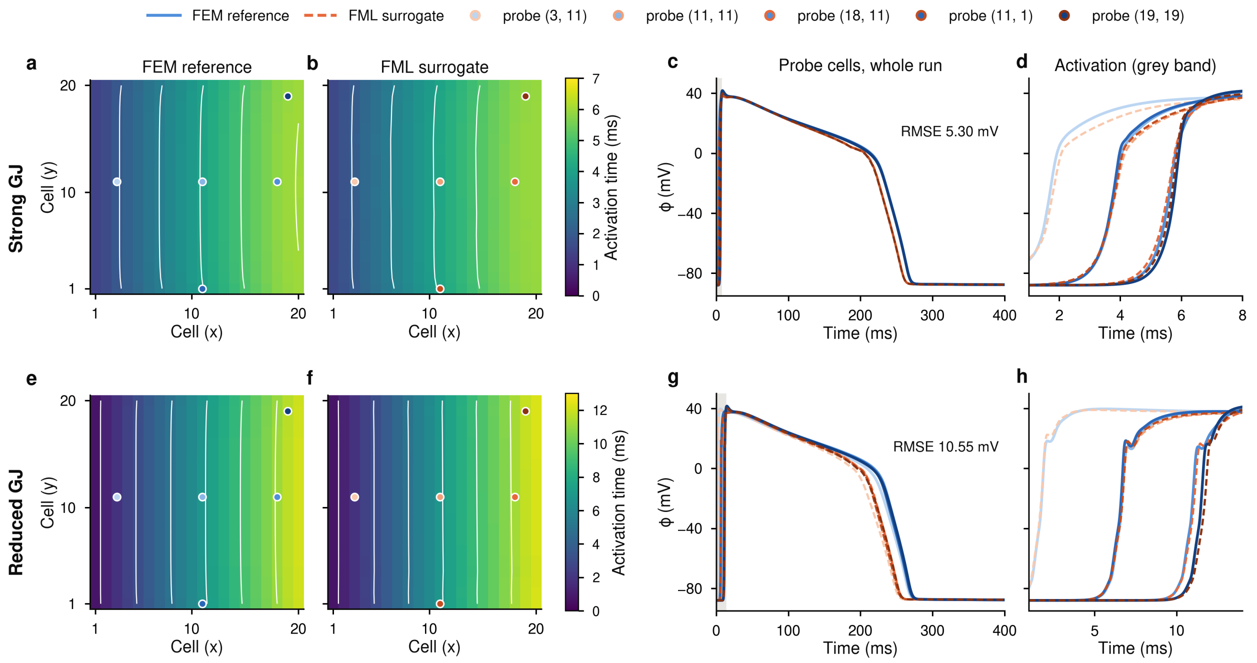


Figure S4: Surrogate against the FEM ID model in a 20 × 20 sheet with mesh 1 at every junction, paced at the left column for one beat of 400 ms. (a–d) Strong and (e–h) reduced gap-junctional coupling. (a,b,e,f) Activation time of every cell in the FEM ID model (a,e) and in the surrogate (b,f), with isochrones (white lines) and the five probe cells (dots). (c,g) Membrane voltage of the probe cells over the run for the FEM ID model (solid) and the surrogate (dashed), with the RMSE of the potential over all 400 cells. (d,h) Upstrokes in the gray band of (c,g).

cycle and core. A fused multiply-add counts as two operations. A Zen 5 core, with two 512-bit FMA units, performs 32 (FP64) and 64 (FP32) per cycle. A CUDA core performs 2 in FP32 and 1/64 of this in FP64. The clock is the all-core base clock for the CPU and the rated boost clock for the GPU. The measured rate is the best of repeated runs of a dense matrix product with the libraries used in the benchmark. The matrices are 8192 × 8192, or 4096 × 4096 for JAX on the CPU and for PyTorch in FP64. Under this load, the GPU runs at 2.88 GHz, above its rated boost clock. At this clock, its FP64 peak is 0.97 TFLOPS, above the 0.96 TFLOPS measured with JAX.

#### Supplementary videos

**Supplementary Video 1: Conduction block in 2-D sheet tissue with regional heterogeneity.** Surrogate simulation of the 100 × 100 sheet of Figure 6a–c, paced at the left edge with the cycle length stepped down from 250 to 206 ms. The central disk of radius 30 cells uses mesh 4 and the rest mesh 1, with reduced gap-junctional coupling and the explicit scheme. Left, membrane voltage of the sheet, with the edge of the disk as a black line. Right, membrane voltage at the four probes, with the current time marked. One frame per millisecond, 30 ms of simulated time per second of video.

**Supplementary Video 2: Conduction block in 3-D block tissue with regional heterogeneity.** Surrogate simulation of the 100 × 100 × 40 block of Figure 6d–f, paced at the left face with the cycle length stepped down from 350 to 205 ms. The central ellipsoid uses mesh 4 and

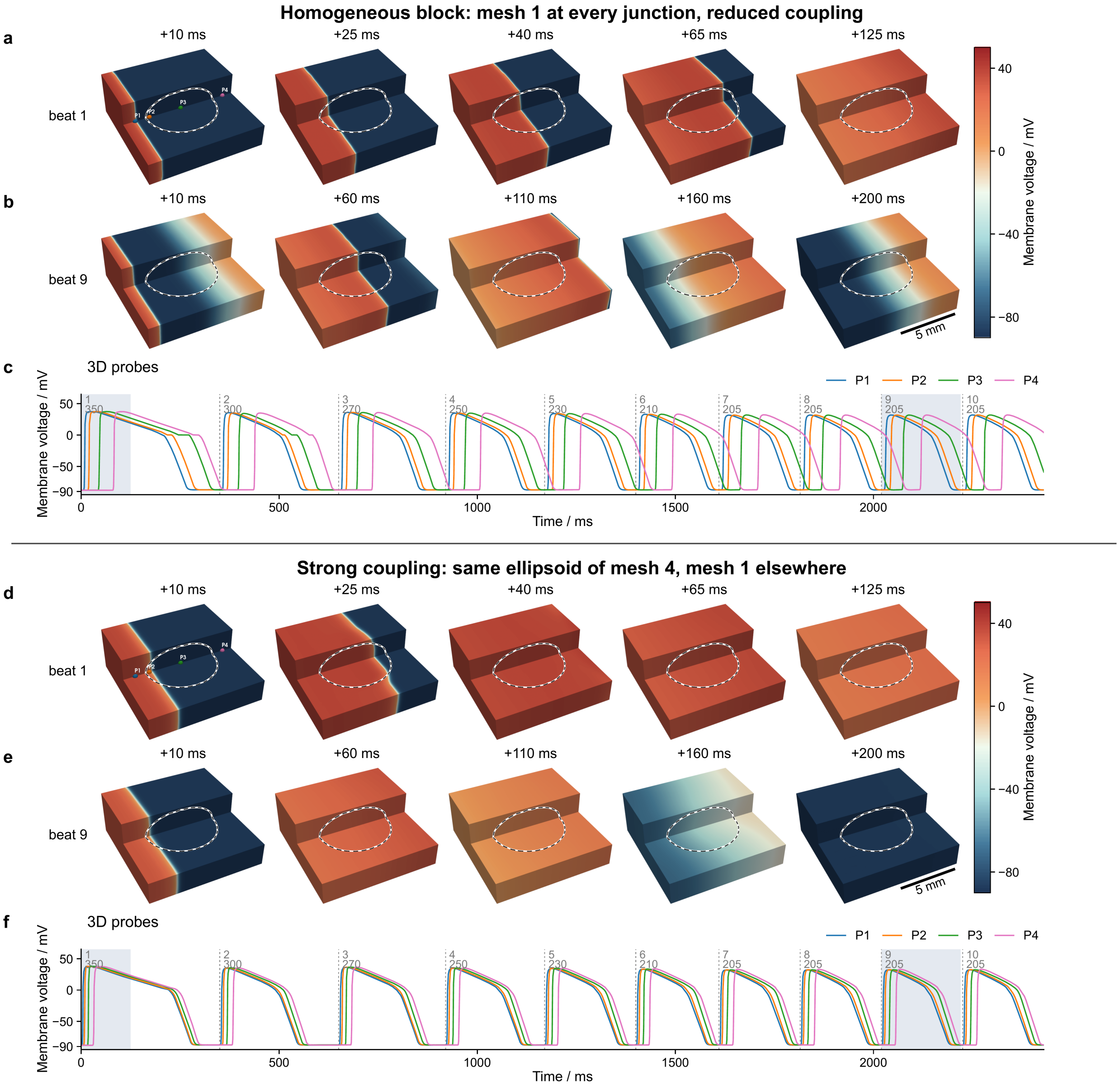


Figure S5: Control simulations of the block of Figure 6d–f, with the same pacing, in which the cycle length steps down from 350 to 205 ms, and the explicit scheme. (a–c) Homogeneous block with mesh 1 at every junction and reduced gap-junctional coupling. The black line marks where the ellipsoid of Figure 6 lies. (d–f) Strong gap-junctional coupling, with the same ellipsoid of mesh 4 (black line) and mesh 1 elsewhere. (a,d) Cut-away views, with the front upper quarter removed, after the first stimulus and (b,e) after the ninth, at the times of Figure 6d,e. (c,f) Membrane voltage at the probes P1–P4 of Figure 6d. Dashed lines mark the stimuli, labeled with the beat number and the cycle length in ms. The shaded windows correspond to (a,b) and (d,e). Scale bars, 50 cells or 5 mm.

Table S2: Hardware and software of the benchmark of Table 1. Floating-point rates in $10^{12}$ operations per second (TFLOPS).

| | |
|---|---|
| CPU | AMD Ryzen 9 9950X (Zen 5, 16 cores / 32 threads, base clock 4.3 GHz, boost up to 5.7 GHz, AVX-512) |
| peak | FP64 2.20, FP32 4.40 TFLOPS ($16 \times 4.3$ GHz $\times$ 32 and $\times$64) |
| measured | FP64 0.87 (MATLAB, Intel MKL 2025.3) and 0.97 (JAX); FP32 2.90 TFLOPS (JAX) |
| Memory | 31 GB usable ($2 \times 16$ GB DDR5-4800, two channels, 76.8 GB $s^{-1}$) |
| GPU | NVIDIA GeForce RTX 5080 (Blackwell, 84 SMs, 10 752 CUDA cores, boost clock 2.62 GHz, 16 GB GDDR7, 960 GB $s^{-1}$), driver 32.0.16.1088 (Windows) / 610.88 (WSL2) |
| peak | FP64 0.88, FP32 56.3 TFLOPS (10 752 $\times$ 2.62 GHz $\times$ 2, FP64 at 1/64) |
| measured | FP64 0.96 (JAX) and 0.85 (PyTorch); FP32 39.1 (JAX) and 40.1 TFLOPS (PyTorch) |
| Operating system | Windows 11 Enterprise (MATLAB, JAX on the CPU, CUDA engine); WSL2 Ubuntu 24.04 (JAX on the GPU) |
| MATLAB | 26.1.0.3346908 (R2026a) Update 5, 16 computational threads |
| JAX | JAX 0.11.2; Python 3.14.0 on Windows (CPU), Python 3.12.3 in WSL2 (GPU, cuda:0) |
| CUDA implementation | PyTorch 2.9.0 with CUDA 12.8, kernels compiled at run time with NVRTC (Windows) |

the rest mesh 1, with reduced gap-junctional coupling and the explicit scheme. The video covers beats 7–10, from 1610 to 2430 ms. Top left, membrane voltage on a cut-away view with the front upper quarter removed and the edge of the ellipsoid as a black line. Top right, the −40 mV surface, that is, the wavefronts and wavebacks, with the ellipsoid in gray. Bottom, membrane voltage at six probes: before the region, in its proximal part, at its center, in its distal part, in a side corridor around it and in the thin layer above it. One frame per millisecond, 50 ms of simulated time per second of video.
